# Serial synapse formation through filopodial competition for synaptic seeding factors

**DOI:** 10.1101/506378

**Authors:** M. Neset Özel, Abhishek Kulkarni, Amr Hasan, Josephine Brummer, Marian Moldenhauer, Ilsa-Maria Daumann, Heike Wolfenberg, Vincent J. Dercksen, F. Ridvan Kiral, Martin Weiser, Steffen Prohaska, Max von Kleist, P. Robin Hiesinger

## Abstract

Following axon pathfinding, growth cones transition from stochastic filopodial exploration to the formation of a limited number of synapses. How the interplay of filopodia and synapse assembly ensures robust connectivity in the brain has remained a challenging problem. Here, we developed a new 4D analysis method for filopodial dynamics and a data-driven computational model of synapse formation for R7 photoreceptor axons in developing *Drosophila* brains. Our live data support a ‘serial synapse formation’ model, where at any time point only a single ‘synaptogenic’ filopodium suppresses the synaptic competence of other filopodia through competition for synaptic seeding factors. Loss of the synaptic seeding factors Syd-1 and Liprin-α leads to a loss of this suppression, filopodial destabilization and reduced synapse formation, which is sufficient to cause the destabilization of entire axon terminals. Our model provides a filopodial ‘winner-takes-all’ mechanism that ensures the formation of an appropriate number of synapses.

## Introduction

Neural circuit assembly requires axonal and dendritic growth followed by synapse formation between specific partners. After pathfinding, axonal growth cones transition to become terminal structures with presynaptic active zones. How axon terminals form a defined number of synaptic contacts with a specific subset of partners is a particularly daunting problem in dense brain regions^1,2^. Stochastically extending and retracting filopodial extensions occur during both pathfinding^3,4^ and synapse formation^5^ and are thought to facilitate interactions between pre- and post-synaptic partners^6-8^. However, very little is known about the role of stochastic filopodial dynamics for robust synapse formation, and how co-regulation of filopodial dynamics and synapse formation is molecularly implemented.

Presynaptic active zone assembly is a key step in synapse formation and regulated by a conserved set of proteins^9,10^. An early active zone ‘seeding’ step has been defined through the functions of the multidomain scaffold proteins Syd-1 and Liprin-α in *C. elegans* and *Drosophila* NMJ^11,12^. Syd-1 is a RhoGAP-domain containing protein^13^ that recruits Liprin-α to the active zone^12^. Liprin-α is an adaptor protein named after its direct interaction with the receptor tyrosine phosphatase LAR (Leukocyte common antigen-related)^14,15^. The Liprin-α/LAR interaction has been directly implicated in active zone assembly across species^16,17^. Downstream of these early factors, Liprin-α and Syd-1 recruit core active zone components and ELKS/CAST family protein Brp^11,18^. Finally, the RhoGEF Trio has been proposed to function downstream of the Lar/Liprin-α/Syd-1^19-21^ and has recently been suggested to regulate active zone size^22^.

Remarkably, the proposed Lar/Liprin-α/Syd-1/Trio pathway has been characterized in parallel for its role in axon guidance, independent of active zone assembly^23-25^. In the *Drosophila* visual system, mutants in all four genes have been implicated in the layer-specific targeting of photoreceptor R7 axons in the medulla neuropil^21,26-30^. It is unclear, whether any of the four mutants affect active zone assembly in R7 neurons. Dual roles in axon pathfinding and synapse formation have been shown or proposed for all four genes^20,21,27,31,32^. Independent implications in active zone assembly and axon pathfinding raise the question whether the two functions reflect independent utilizations of a pathway ‘module’ in different contexts, or whether the two functions are in fact connected.

In this study, we investigated the relationship between filopodial dynamics and synapse assembly in the presynaptic R7 terminal. We identified a rare type of filopodium that only occurs during the time of synapse formation. The early synaptic seeding factors Liprin-α/Syd-1 accumulate in only a single such filopodium per terminal at any given point in time. Consequently, only one such filopodium per terminal is stabilized, suggesting that only one filopodium is competent to form a synapse at a given time point. A data-driven computational model shows that this ‘serial synapse formation model’ is supported by the measured dynamics and could be tested in mutants. Correspondingly, loss of *liprin-α* or *syd-1* specifically affects the stability of these rare filopodia, while loss of *lar* affects their formation and *trio* selectively abolishes the negative feedback on their formation. These highly specific filopodial effects precede all other defects in these mutants, including axon terminal retractions. Our findings support a ‘winner-takes-all’ mechanism in which filopodial competition ensures synapse formation in a serial manner through non-random distribution of early synaptic seeding factors. We provide a quantitative model from stochastic filopodial dynamics to the formation of a limited number of synapses as well as a model for axon terminal stabilization based on filopodia and synapses.

## Results

To understand the role of filopodial dynamics during synapse formation in the context of normal brain development we chose *Drosophila* R7 photoreceptors as a model. In each optic lobe, ~800 R7 axon terminals reach their adult morphology as column-restricted, smooth, and bouton-like structures that contain around 20-25 presynaptic release sites (**Fig. 1a-b**)^33, 34^. In contrast to the smooth adult structure, during synapse formation these axon terminals exhibit highly dynamic filopodial extensions (**Fig. 1a**). While R7 filopodial dynamics during the first half of pupal development (P+0-50%) are thought to play a role in axon guidance and layer formation^5^, the role of filopodial dynamics in the second half of pupal development (P+50-100%) is unknown.

**Figure 1:**
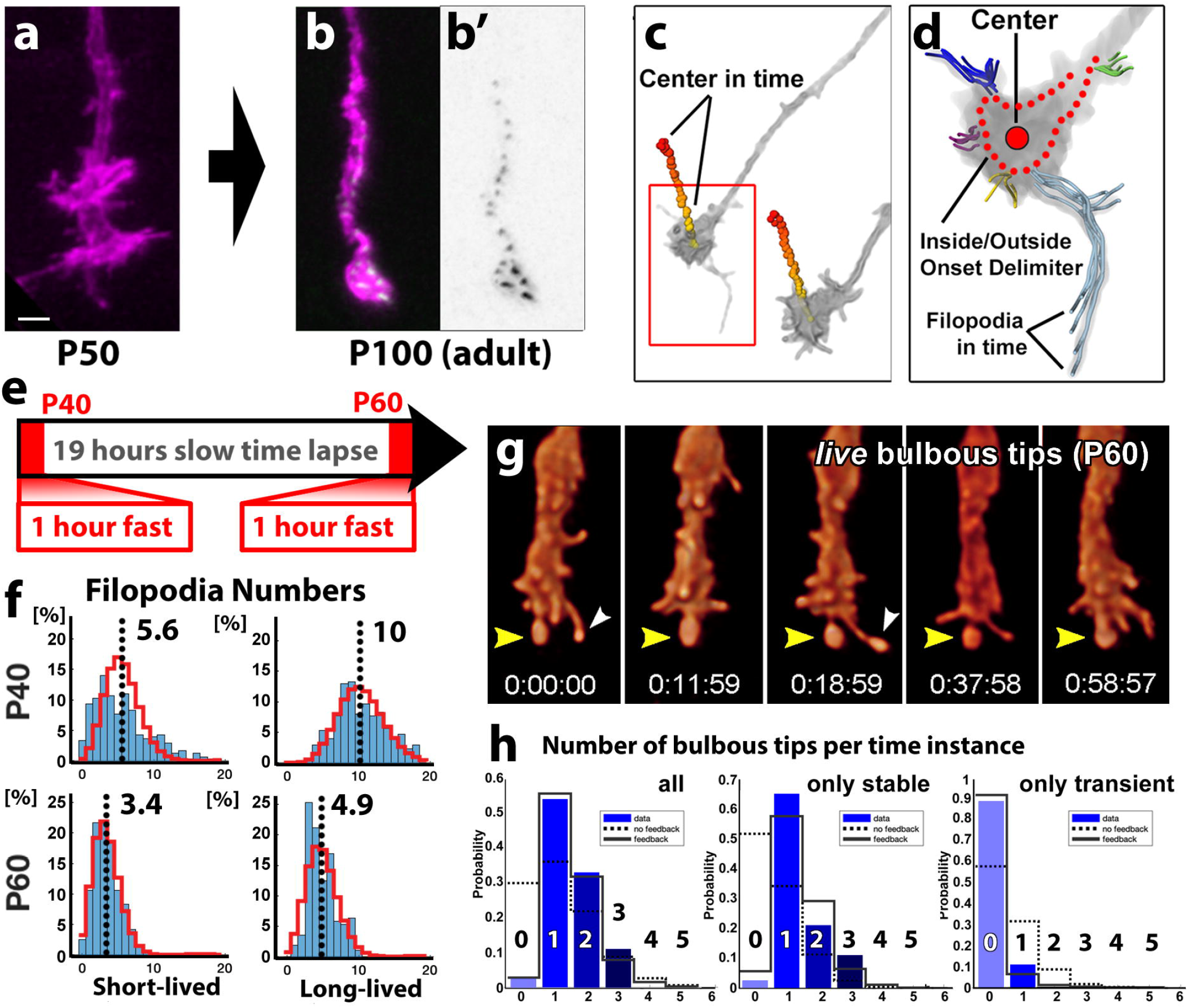
4D filopodia tracking reveals stochastic dynamics prior to synapse formation and rare ‘bulbous’ filopodia that stabilize one at a time during synapse formation. (a, b) *Drosophila* R7 photoreceptor axon terminals transition from a growth cone-like structure with multiple filopodia just prior to synapse formation at 50% pupal development (P50) to a smooth, adult terminal with 20-25 synapses (magenta: CD4-tdTomato, green: BrpD3-GFP). (c, d) A semi-automatic method for 4D filopodia tracking is based on the Amira filament editor. (e) Imaging protocol for >20 hour continuous time lapse and fast imaging at P40 and P60 for the same axon terminals. (f) R7 filopodia fall into short-lived and long-lived classes that both fit Poisson (stochastic) distributions at both P40 and P60. (g) Representative snapshots of an R7 axon terminal in the brain at P60 with a continuous stable bulb (yellow arrowhead). (h) Number of bulbous filopodia at P60. Separation into stable (middle) and transient (right) bulbs reveals that most time points contain 1-2 stable bulbs. The distributions can be fit with negative feedback (sold black lines), but not with Poisson product distributions (dotted lines). Scale bar in (a) for (a, b): 2um.

### 4D filopodia tracking reveals stochastic dynamics prior to synapse formation and rare ‘bulbous’ filopodia that stabilize one at a time during synapse formation

The characterization of axonal filopodia dynamics during synapse formation in the intact brain required a method to obtain quantitative high-resolution 4D data throughout the second half of fly brain development. We have previously developed long-term culture of intact developing brains in an imaging chamber^5^, but difficulties in tracking fast dynamics for thousands of filopodia in a developing brain have so far precluded large-scale analyses. We therefore devised a semi-automatic method for standardized and quantitative filopodia tracking based on a previously developed ‘filament editor’ (**Suppl. Fig. 1a-b**)^35^. In short, the newly developed algorithm predicts the growth cone centers for all time points by similarity-based propagation from the initial time point and thereby streamlines the segmentation of individual terminals (**Fig. 1c**). Next, filopodia are traced at each time point, sequentially propagated, and automatically matched to the corresponding filopodia at other time points based on the vicinity of their starting points (**Fig.1d**). Using this method, we tracked 27,390 individual filopodia through time and space across 38 growth cones for this study (Methods, **Suppl. Fig. 1a-b**).

We first analyzed 4D datasets of wild-type R7 axon development from just before synapse formation (P+40%, P40) until after 20-22h in culture, during synapse formation (P+60%, P60) for the same growth cones (spatial resolution: voxel size 0.1×0.1×0.5 μm; temporal resolution: 1 minute time lapse for 1 hour periods; **Fig. 1e**). 4D tracking of several thousand filopodia revealed two distinct classes with separate exponential lifetime distributions: highly transient filopodia with a maximum lifetime of 8 minutes (short-lived), and stable filopodia with lifetimes of more than 8 minutes (long-lived) (**Suppl. Fig. 1c**) with similar length and velocity distributions (**Suppl. Fig. 1d**). At any time instance at P40 each R7 terminal has twice as many long-lived (>8 min) compared to short-lived (<8 min) filopodia (**Fig. 1f**). The numbers of both classes of filopodia reduce significantly by P60 (**Fig. 1f**), and by P100 all filopodia disappear (**Fig. 1b**). The measured filopodia exhibit linear stochastic dynamics, since all four distributions (numbers of long- and short-lived filopodia at P40 and P60) almost perfectly fit Poisson distributions^36^ (red traces in **Fig. 1f**; Suppl. Note on Mathematical Modeling).

In addition to the great majority of transient filopodia, we also consistently observed rare long-lived filopodia that develop characteristic ‘bulbous tips’ around the time of synapse formation^5^. Quantitative analysis revealed no bulbous tips prior to synapse formation at P40. In contrast, at P60 there are 1-2 stabilized filopodia with bulbous tips present at any time point, most of which has a lifetime of >40 minutes during the acquisition window of 1-hour used for fast dynamics (**Fig. 1g-h; Suppl. Movie 1**). Because many of these filopodia existed before and after the 1-hour imaging window, the lifetime estimate is certainly an underestimation, and we observed bulbous tips that existed for hours in long-term time lapse. Notably, we counted at most time instances only one or two bulbous tip filopodia (bulbs) per axon terminal at any given time point, and almost no time instances with 0 bulbs (**Fig. 1g-h; Suppl. Movie 1**). This heavily right-skewed distribution is indicative of a regulatory mechanism: While the absence of regulatory mechanisms would give rise to a Poisson product distribution (dotted lines in **Fig. 1h**), the inclusion of an inhibitory feedback, whereby existing bulbs suppress new bulbs, reveals an excellent fit of the observed distribution (solid lines in **Fig. 1h**). This skewed distribution causes a bulbous tip to be present at almost every time point, while a Poisson product distribution (= no feedback) would result in many time points without bulbs. Correspondingly, transient bulbs occur comparatively rarely, while 65% of all time points have exactly one bulb (**Fig. 1h**) and almost 100% of time points have *at least* one bulb. Hence, at the time of synapse formation, R7 growth cones continuously stabilize a single filopodium at any given time point, while the overall number of filopodia decreases.

### One filopodium at a time accumulates synaptic seeding factors

Competitive destabilization of secondary bulbous filopodia could be achieved through a ‘winner takes all’ mechanism of filopodial competition. We asked whether synaptic building proteins would exhibit such a competitive distribution at filopodial tips. First, we tested whether the active zone protein Burchpilot (Brp) is associated with filopodia. Fluorescently tagged BrpD3 (or Brp^short^) is a reliable marker for mature synapses and localizes specifically to sites of intrinsic Brp without affecting synapse function or causing overexpression artefacts seen with other genetically encoded synaptic markers, including full-length Brp-GFP^37-39^. However, we never found Brp-marked mature active zones in filopodia, similar to recent findings in developing adult motoneurons^40^. (**Fig. 2a-b, g**). To measure the dynamics of synapse formation, we performed live imaging of BrpD3 at 10 min resolution over several hours around P+70% (**Suppl. Movie 2**). Brp signal above noise levels was strikingly excluded from filopodia; puncta never moved into or formed in filopodial tips, and instead formed by gradual accumulation on the axon terminal main body as it transitions into the smooth terminal bouton of the adult. Tracking individual puncta for over 5 hours revealed that the vast majority of Brp-positive synapses are stable once formed (**Fig. 2h, Suppl. Movie 2**). We conclude that mature synapses marked by Brp are not associated with filopodia, but only form on the axonal trunk where they are stable once formed.

**Figure 2:**
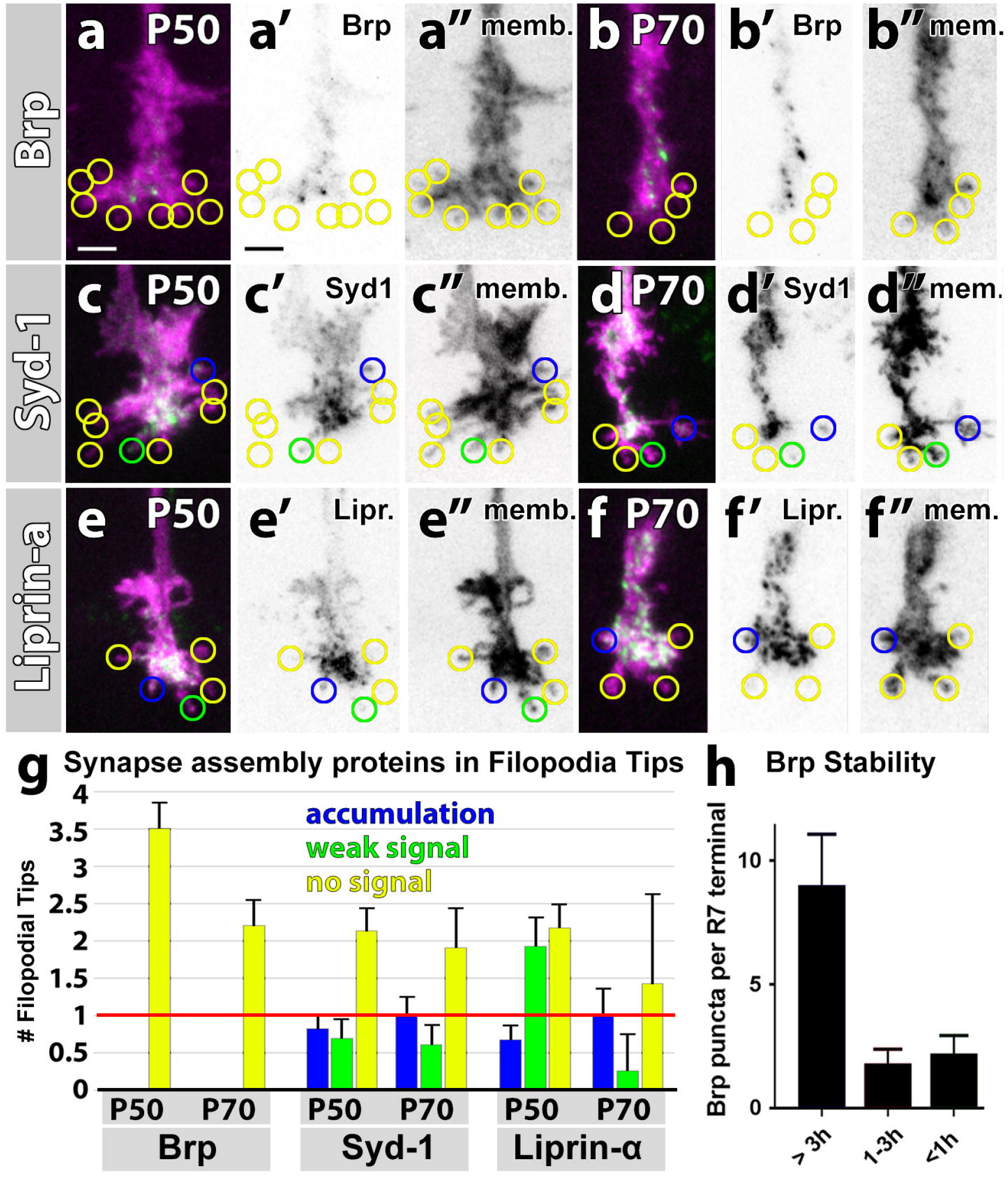
One filopodium at a time accumulates synaptic seeding factors. (a-f) Localization in R7 photoreceptor terminals and filopodia for BrpD3-GFP (a, b), GFP-Syd-1 (c, d) and Liprin-α-GFP (e, f). Shown are two time points: P50 (a, c, e) and P70 (b, d, f). Yellow circles indicate filopodia with no measurable GFP signal, green circles weak signal, and blue circles clear accumulations. (a’-f’) show the single channel for the GFP-tagged proteins (green), and (a”-f”) show the single channel for the membrane tag CD4-tdTomato, which marks all filopodia (magenta). Scale bar: 2 μm. (g) Quantification of filopodial accumulation of the three proteins. (h) Number of BrpD3 punctae per R7 terminal binned according to their lifetimes. R7 terminals were live imaged at 10 min resolution starting at P+50% + 22h in culture. Individual punctae were tracked for 5,5h to determine lifetimes (n = 5 terminals). Error bars denote SEM.

At the *Drosophila* neuromuscular junction, Brp is recruited late to nascent synapses by the early seeding factors Syd-1 and Liprin-α^12^. Antibodies against either protein labeled a high density of puncta throughout the brain that did not allow to assign localization to individual filopodia. We therefore overexpressed GFP-tagged variants of either protein and asked where an overabundance of Liprin-α and Syd-1 would localize. Unlike Brp, GFP-tagged Liprin-α and Syd-1 occurred in bulbous filopodia tips (**Fig. 2c-g**). Remarkably, clearly discernable accumulations of Liprin-α and Syd-1 were only apparent in one or sometimes two bulbous filopodia, while the majority of filopodia contained no signal (**Fig. 2c-g**). In contrast to other filopodia, the number of filopodia tips containing Syd-1 or Liprin-α did not decrease between P50 and P70 but remained constant at 1 per axon terminal. Note that most filopodia did not contain any detectable Syd-1 or Liprin-α, despite large amounts of overexpressed proteins in the axon terminal trunks. The single positive filopodia could not be predicted based on size or length of the filopodia (compare c’-f’ to c”-f”). Hence, Syd-1 and Liprin-α match the criteria for a ‘winner takes all’ distribution: both localize sparsely and non-randomly to, on average, one single filopodium per axon terminus at both P50 and P70.

Live imaging revealed the localization of Liprin-α-GFP only to stable filopodia with bulbous tips, while dynamically moving in and out of the filopodium (**Suppl. Movie 3**); GFP-Syd1 puncta were too dense for reliable tracking, but similar to Liprin-α, only exhibited clear accumulations in bulbous tips. These observations suggest that synapse assembly may start in filopodia. The data are further consistent with reversible molecular ‘seeding’ events^12^ and filopodia stabilization through nascent synapses, as previously observed^40,41^. The findings suggest a model whereby only one filopodium at a time is stabilized and contains an increased amount of synaptic seeding proteins Syd-1 and Liprin-α. In turn, only one filopodium at a time may be synaptogenic, i.e. competent to form a synapse. This ‘serial synapse formation’ model would provide a regulatory mechanism for the generation of a limited number of synapses throughout the second half of pupation. Furthermore, a feedback model based on a recruitment of seeding factors to stabilized bulbous filopodia would rely on continuous exchange of material across filopodia, suggesting a need for continuous dynamics.

### A data-driven computational model predicts ‘serial synapse formation’ based on competition and negative feedback of bulbous filopodia

To test whether the ‘serial synapse formation’ hypothesis is consistent with the measured dynamics, we built a data-driven Markov jump model of filopodia dynamics based on the measured data at P40 and P60 (see also Suppl. Notes on Mathematical Modeling). Specifically, we modeled the birth of filopodia, their transitions between short-lived and long-lived filopodia, transitions to bulbous filopodia, and finally transitions to synapses (**Fig. 3a**). To account for the ‘winner takes all’ bulb distributions we included inhibitory feedback based on the measured bulb distribution and the filopodia competition model. The live imaging data provided direct measurements of the filopodial birth and death rates (r1 and r2) and the observed rates of bulb disappearance (r4) (All measured data are labeled in blue). Because of the introduction of inhibitory feedback on bulb formation, the average rate of bulb initiation (r3) at P60 is the product of a propensity to form bulbs (r2B) and the average inhibitory feedback f1, such that absence of feedback (f1=1) represents no inhibition of r3 and maximal negative feedback (f1=0) represents complete inhibition of r3. Based on the measurement of bulb appearances and the observed right-skewed distribution of bulbs (**Fig. 1h**) we determined the exclusive set of r2B and f1 that fit the observations for time point P60 (**Suppl. Table 1**). Negative feedback f1 in WT is close to maximal, which ensures the measured sharp distribution of only 1-2 bulbs per time instance with almost no time instance of zero bulbs. As shown in **Figure 1h**, almost every transient bulb stabilizes (r5). Lastly, we estimated the rate of synapse formation (r6) from the maximal slope in **Fig 3f**. As a cross validation of the model, we observe that the number of mature synapses matches previous measurements^33,34^.

**Figure 3:**
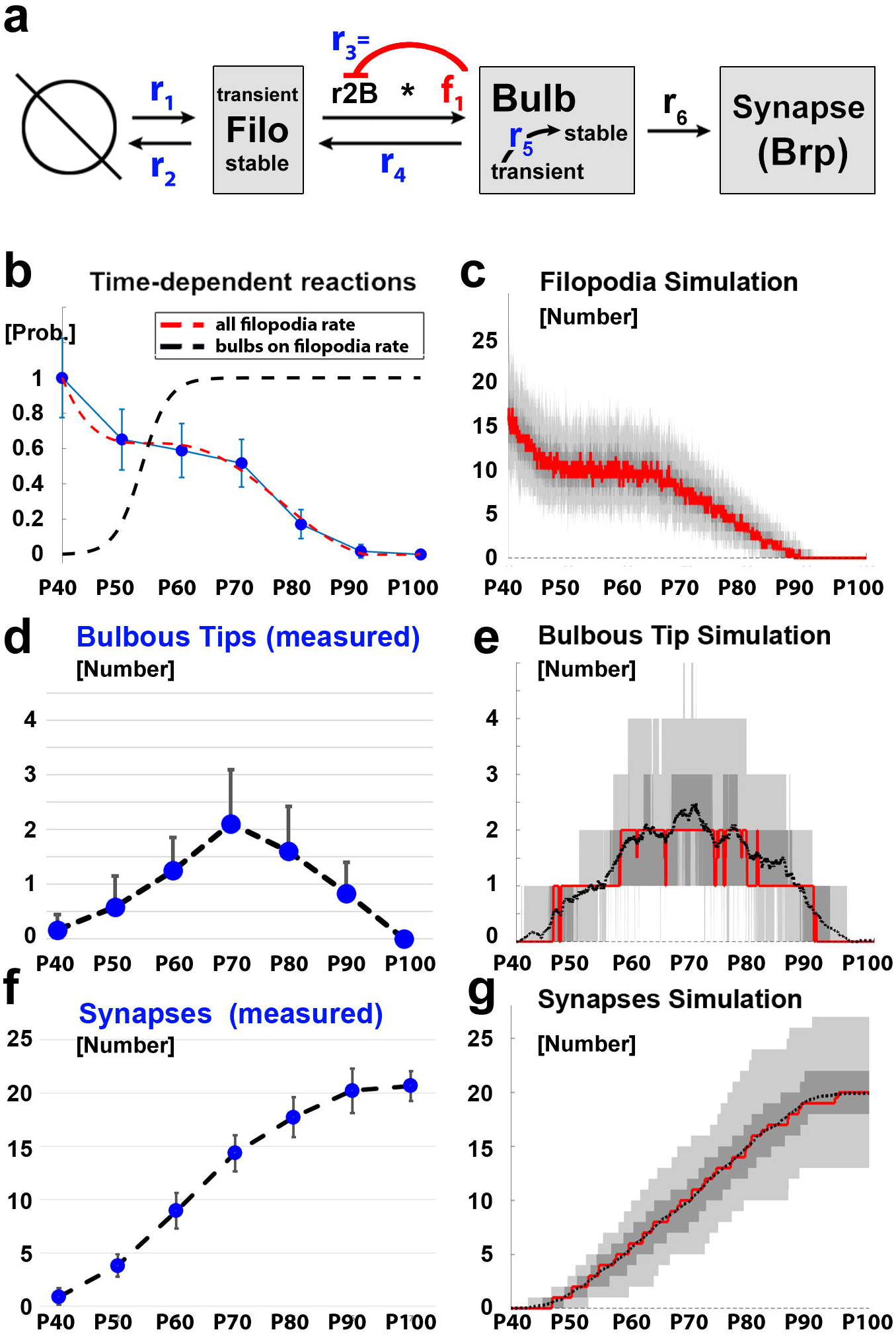
A data-driven computational model predicts ‘serial synapse formation’ based on competition and negative feedback of bulbous filopodia. (a) Summary of the data-driven Markov state model from filopodial birth to synapse formation. All rates in blue are measured from live imaging data. (b) Estimation of time-dependent function required for the modeling from P40-P100 (40%-100% of pupal development). The reduction of filopodia was based on measured filopodial counts from fixed preparations (blue disks). The increased propensity to form bulbs on these filopodia was estimated based on bulb measurements shown in panel (d). (c) Output of Markov state model for filopodial dynamics based on measured rates according to the model in (a). (d) Measured number of bulbous tips. (e) Output of Markov state model for the development of bulbous tips. Black dotted lines: mean number of bulbs; solid red line: median number of bulbs; dark grey denotes the interquartile range (50% of the data) and light grey the 95% confidence range. (f) Measured numbers of synapses between P40 and P100. (g) Output of Markov state model for synapse formation.

In addition to the live dynamics measurements at P60, we counted total numbers of filopodia, bulbous tips and synapses (BrpD3) in fixed preparations for the time points P40, P50, P60, P70, P80, P90 and P100 (blue data points in **Fig. 3b, d, f**). The live P60 data matched the fixed counts at P60 well. Based on these data, we determined a function for the filopodial decline (red dotted line in **Fig. 3b**) and the propensity to form bulbs (black dotted line in **Fig. 3b**). Based on the measured data and these two rates, we modeled the changes to types (Filopodia and Bulbs) and numbers of filopodia over time in 3600 time steps, equivalent to 3600 minutes from P40 to P100. The resulting model reproduces a minute-by-minute simulation of the number of filopodia (**Fig. 3c**), bulbs (**Fig. 3e**) and synapses (**Fig. 3g**).

The appearance of only 1-2 bulbous tips at any time point between P55 and P85 leads to a continuous, limited generation of mature synapses that matches well with measured BrpD3 data (**Fig. 3f, g**). Furthermore, the model predicts variability of synapse numbers similar to the measured variability. For model building and parameter estimation based on the measured data, see Suppl. Note on Mathematical Modeling. We conclude that our serial synapse formation model, based on measurements of filopodia and competitive feedback between bulbs, can in principle explain the kinetics and distribution of synapse development observed in wild type.

### Loss of synaptic seeding factors Syd-1 and Liprin-α causes a loss of inhibitory feedback during filopodial bulb formation

Our model requires filopodial competition that stabilizes only one bulbous filopodium at a time. Similarly, Syd-1 and Liprin-α accumulate in one filopodium at a time, while the late synaptic assembly protein Brp does not localize to filopodia (**Fig. 2**). We therefore hypothesized a role for the synaptic seeding factors in bulb stabilization and tested our model experimentally in mutants for these factors. First, we asked whether filopodia and growth cone dynamics depend on Brp. We tested the consequences of a loss of *brp* during R7 axonal development with a previously tested combination of two RNAi constructs^42^ and confirmed the known defect in neurotransmission (**Suppl. Fig. 2a-b**). The knock-down of Brp had no effect on the transition of R7 terminal morphology from filopodial to smooth bouton-like structures (**Suppl. Fig. 2c**). These findings indicate no role for Brp in axon terminal development and are consistent with the absence of Brp from filopodia (**Fig. 2**). These findings further resemble previous observations in motoneurons^40^, and are consistent with the observation of normal development in the absence of neurotransmission^43^.

To perturb early stages of synapse formation, we investigated mutants for *liprin-α* and *syd-1.* The analysis of filopodial dynamics is complicated by previous observations of R7 axon targeting defects for both *liprin-α* and *syd-1*^21,28,29^. To characterize the timeline and origin of these defects, we performed long-term live imaging from P+30%-P+70% for single mutant, positively labeled R7 cells in an otherwise heterozygous background (MARCM)^44^. Our analyses of both mutants (*liprin-α^E^^28^, syd-1*^w46 21^) revealed that in both cases all mutant R7 axons initially targeted correctly. Axon terminal dynamics of both liprin-*α* and *syd-1* mutant axons are indistinguishable from wild type until P40 (**Suppl. Fig. 3a-c**). Starting around P50, i.e. during the time period of synapse formation, individual terminals retract. At P60, the majority of *liprin-α* or *syd-1* mutant R7 axon terminals continue to remain in their correct target layer. Our long-term live imaging datasets therefore allow to distinguish and separately analyze filopodial dynamics of terminals that remain stable while synapse formation commences from P40 - P60 and identify those terminals that will retract later during development. We therefore first performed a quantitative analysis of filopodial dynamics at P40 and P60 for those terminals that remained stable in their correct target layer and are therefore independent of, or preceding, retraction. We provide a detailed analysis of retraction events in the last section (**Fig. 6**).

Quantitative 4D tracking of filopodia of both *liprin-α* or *syd-1* mutant R7 axon terminals at P40 and P60 revealed distributions of numbers, lifetimes, lengths and velocities that are largely indistinguishable from wild type (**Suppl. Fig. 3a-o**). *syd-1* mutant axon terminals exhibited individual, unusually elongated filopodia during synapse formation, but their low number did not affect the statistics significantly (**Suppl. Fig. 3m**).

In contrast to short- and long-lived filopodia, the dynamics of bulbs were significantly altered. The lifetimes of bulbous tips in both *syd-1* and *liprin-α* were reduced by 70-80% (**Fig. 4a**). Bulbs destabilization is also reflected in a similar ~70% decrease of the number of stable (>40 minute) bulbous filopodia in both mutants. Correspondingly, the number of short-lived, destabilized bulbs is dramatically increased by ten to twenty-fold (**Fig. 4b, Suppl. Movie 4**). The reduced lifetimes and increased numbers are a result of increased rates for both bulb appearance (r3) and bulb disappearance (r4) as measured at P60 (**Suppl. Table 1**). A remarkable consequence of corresponding increases in both bulb generation and destabilization is that the average number of bulbs observed per time instance, i.e. the average appearance of what a fixed image would look like, is not significantly different from wild type (**Fig. 4c**). The observed bulb destabilization and corresponding overproduction of transient bulbs suggest a compensatory mechanism. In this case, the absence of synaptic seeding factors would lead to a defect in bulb stabilization, resulting in continuous attempts to form new bulbs. Bulbs stabilization by synaptic seeding factors is consistent with the tightly controlled, non-random distribution of Liprin-*α* and Syd-1 proteins as well as the wild type bulb distributions and further indicates that the machinery for bulb initiation *per se* is robust and independent of Liprin-α or Syd-1. We conclude that synaptic seeding factors are required for bulb stabilization, but not for bulb initiation.

**Figure 4:**
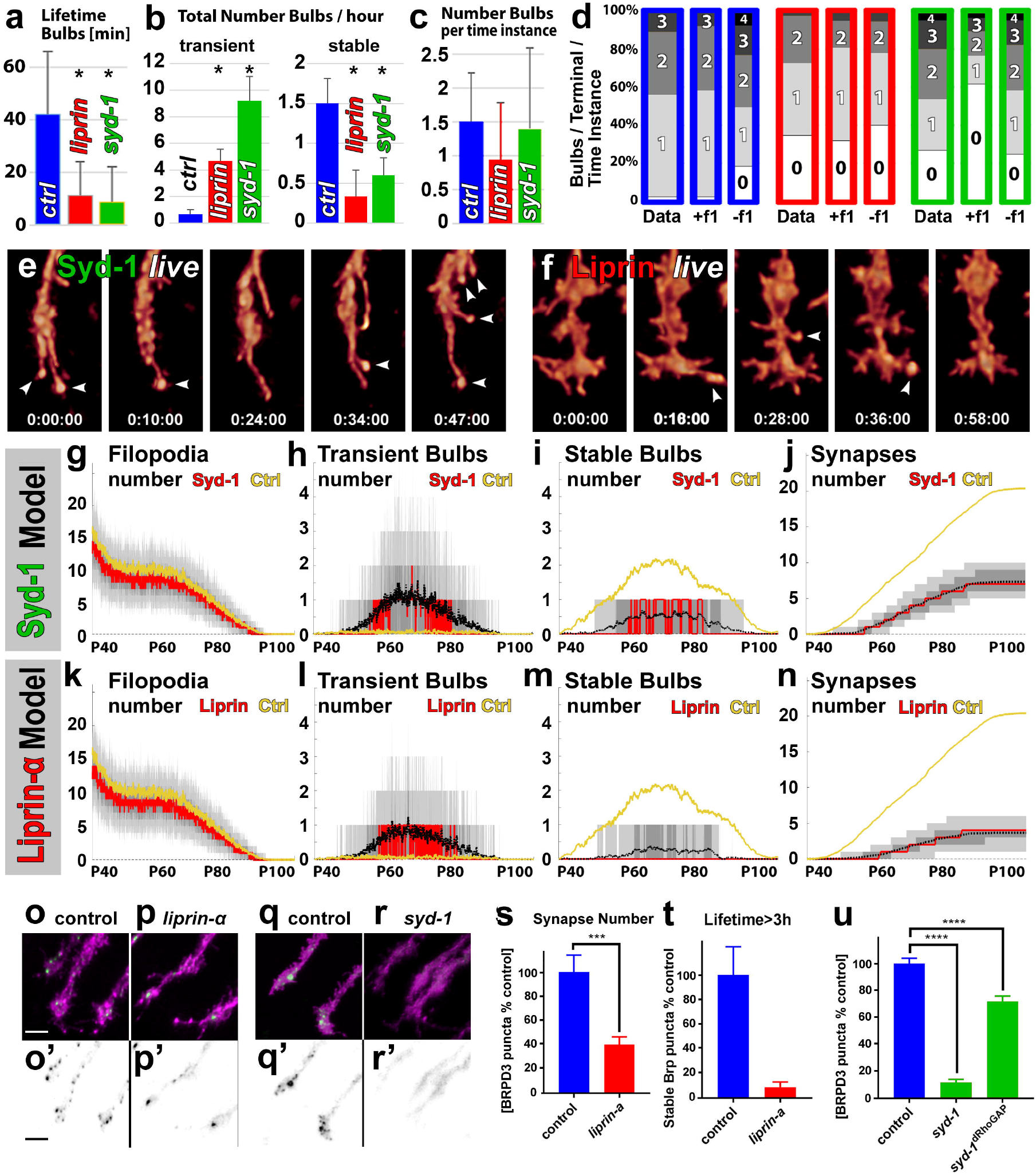
Loss of synaptic seeding factors Syd-1 and Liprin-α causes a loss of inhibitory feedback and filopodial bulb destabilization. Analyses of filopodial dynamics and synapses formation for *syd-1* (green), *liprin-α* (red) and control (blue). (a) Lifetime of bulbous filopodia. (b) Total number of bulbous filopodia per terminal per hour. (c) Average number of bulbous filopodia per time instance. (d) Number of concurrently existing bulbous filopodia per axon terminal per time instance. (e, f) Representative snapshots of *liprin-α* (e) and *syd-1* (f) revealing only transient bulbs. (g-n) Markov state model simulation for *syd-1* (g-j) and *liprin-α* (k-n) for the numbers of filopodia (g, k), transient bulbs (h, l), stable bulbs (i, m) and synapses (j, n). In all cases control traces from Fig. 3 are shown in yellow. Black dotted lines: mean number of bulbs; solid red line: median number of bulbs; dark grey denotes the interquartile range (50% of the data) and light grey the 95% confidence range. (o-r) Measurement of BrpD3 punctae in mutant axon terminals. (o’-r’) BrpD3 single channel. Scale bar: 2 μm. (s, t) Quantification of BrpD3 synapse numbers per terminal relative to control. n=18 and 16 (p = 0.0007). (t) Number of BrpD3 punctae per terminal with lifetimes greater than 3h in R7 axons live imaged for 4h at P+70% in wild-type (n =5) and *liprin-α*^E^ mutants (n=5). (u) Quantification of synapse numbers in viable flies for a precise genomic deletion of the putative RhoGAP domain of *syd-1.* n = 45, 18 and 32 (p < 0.0001). Error bars denote SEM

Next we analyzed the distribution of bulbs present at any given time point over a one hour period at P60. In contrast to wild type, both *syd-1* and *liprin-α* exhibited 20-30% of all time points without any bulbs (**Fig. 4d**). In contrast to the bulb distributions in wild type (blue boxes), both *syd-1* and *liprin-α* (green and red boxes, respectively) resemble Poisson product distributions and are not better matched by applying the inhibitory feedback necessary to fit the wild type data (**Fig. 4d**). As with the wild type data, the observed rate of bulb appearance (r3) could be fitted with a single product of the average propensity to form a bulb (r2B) and the average inhibitory feedback (f1) at P60 (**Fig. 3a**). In both *syd-1* and *liprin-α* the feedback was mostly lost (f1 >10-fold increased, **Suppl. Table 1**). Morphologically, these observations result in a high frequency of transient bulbs for both *liprin-α* (**Fig. 4e; Suppl. Movie 4**) and *syd-1* (**Fig 4f; Suppl. Movie 4**).

We next used our simulation of the entire time course from P40-P100 from filopodial dynamics to synapse formation established for wild type (**Fig. 3**) using the measured live data at P60 for each mutant. For *syd-1,* overall filopodia numbers are slightly below wild type (**Fig. 4g**). However, in contrast to wild type, large numbers of transient bulbs form (**Fig. 4h**), but very few stabilize (**Fig. 4i**). In contrast, wild type forms almost no transient bulbs, because all bulbs stabilize (compare black trace for mean in *syd-1* to yellow traces of the mean for control in **Fig. 4h-i**). The *liprin-α* simulation revealed a similar increase of transient bulbs combined with a further reduced number of stable bulbs (**Fig. 4k-m**). As a result of this altered bulb distribution, the simulation predicts a reduction of adult synapses in *syd-1* and *liprin-α* to 35% and 20% of the wild-type levels, respectively (**Fig. 4j, n**). These simulated reductions occur without changes to the synapse formation rate r6 and purely because of the observed defect in bulb stabilization; an additional direct effect of *syd-1* or *liprin-α* on synapse formation itself, as has been argued based on their molecular function as synaptic seeding factors^12^, would reduce the number of synapses further.

To assess the number of synapses *in vivo* we performed BrpD3-based counts at P70 in R7 axon terminals in their correct target layer. The number of Brp-positive synapses was significantly reduced in *liprin-α* (**Fig. 4o-p, s**) and almost completely abolished when a 3 hour stability criterion is applied (**Suppl. Movie 5, Fig. 4t**). Similarly, *syd-1* mutants exhibited almost no Brp-positive synapses (**Fig. 4q-r**). We also generated a precise CRISPR-mediated knock-in of a mutant version of *syd-1* lacking putative RhoGAP activity, which was previously predicted to play a role in active zone assembly^45 22^ (**Suppl. Fig. 4a-b**). However, in contrast to loss of *syd-1,* the *syd-1^ΔRhoGAP^* mutant flies are viable, fertile and exhibit no obvious defects other than a relatively mild reduction of BrpD3-positive synapse numbers (**Fig. 4u, Suppl. Fig. 4c-d**) and are not further analyzed here. Consistent with studies in other systems, these findings indicate that *syd-1* and *liprin-α* are required for normal synapse formation also in R7. The defects in synapse formation are independent of axonal retractions, because all analyses presented here were done on axon terminals that remained stabely in the correct target layer. Our findings support the hypothesis that both proteins function as a limiting resource for synaptic ‘seeding’, which is in turn required for bulb stabilization. The altered dynamics of bulbous filopodia is sufficient to account for a substantial reduction in synapses.

### Analysis of the Syd-1/Liprin-α pathway components reveal a role for Lar, but not Trio in bulb initiation

The membrane receptor LAR and the RhoGEF Trio have been proposed to function in a pathway with Liprin-α and Syd-1 in both contexts, axon pathfinding and synapse formation^20,21,27,31^. We therefore performed live imaging, filopodia tracking, quantitative analyses and computational modeling for *lar* and *trio* mutant R7 axons analogous to WT, *syd-1* and *liprin-α.*

Long-term live imaging of single *lar* or *trio* mutant R7 axons showed that, similar to *liprin-α* and *syd-1,* both *lar* and *trio* mutant axon terminals initially target correctly and exhibit no significant alterations of their filopodial dynamics prior to P40 (**Suppl. Fig. 5a-c**). However, individual *lar* mutant axon terminals exhibit the first probabilistic retractions shortly thereafter, resulting in retraction of nearly all terminals by P70, as previously reported^26,27^. In contrast, we did not observe any retractions of *trio* mutant axons. As in the *syd-1* and *liprin-α* mutants, we analyzed filopodial dynamics at P40 and P60 exclusively for axon terminals that targeted normally and exhibited stable dynamics in their correct target layer throughout. An analysis of retractions will follow in the last section (**Fig. 6**).

Similar to *liprin-α* and *syd-1,* the dynamics of *lar* mutant R7 growth cones exhibited no significant differences of filopodia numbers, lifetimes and lengths until P40 (**Suppl. Fig. 4a-c**). *trio* mutants exhibited mild increases in numbers based on an increased birth rate r1 (**Suppl. Table 1**). However, both short-lived and long-lived filopodia exhibited distributions for numbers and lengths that were similar to wild type in both mutants (**Suppl. Fig. 4d-o**). In contrast, bulbous tip dynamics were significantly affected in both mutants. Similar to *syd-1* and *liprin-α,* both mutants exhibited bulbs of significantly reduced lifetimes (**Fig. 5a, Suppl. Movie 6**). Also similar to *syd-1* and *liprin-α,* bulbs were destablized in *lar* (**Fig. 5b**). However, in contrast to *syd-1* and *liprin-α, lar* mutants form overall significantly less bulbs, suggesting a defect in bulb formation (**Fig. 5d**). In contrast, *trio* exhibited a strong increase of transient bulbs without significant loss of stable bulbs and a normal average number of bulbs per time instance (**Fig. 5b, c**). Hence, *trio* has both stabilized bulbs as well as a large number of destabilized bulbs.

**Figure 5:**
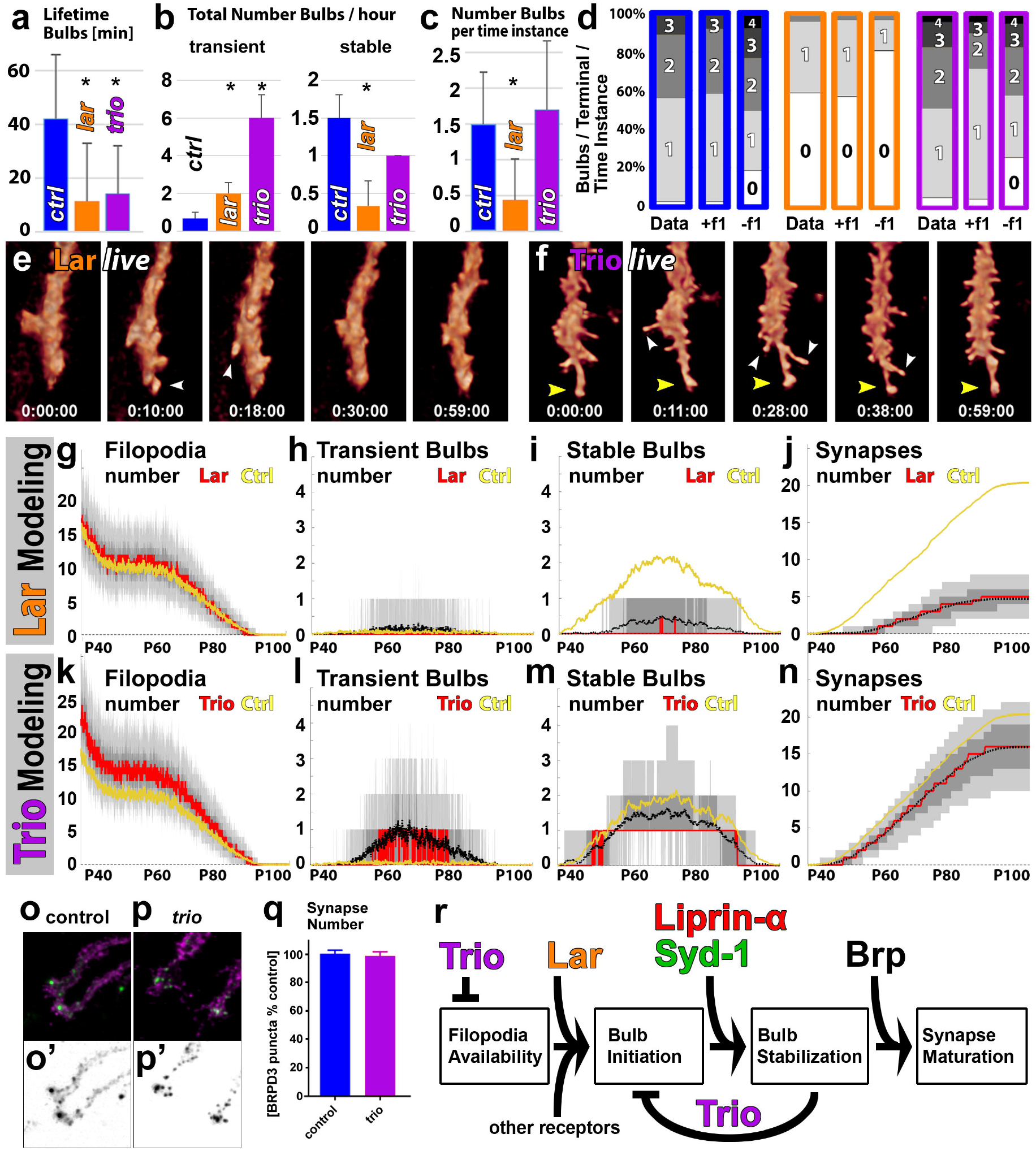
Analysis of the Syd-1/Liprin-α pathway components reveal a role for Lar, but not Trio in bulb initiation. Analyses of filopodial dynamics and synapses formation for *lar* (orange), *trio* (magenta) and control (blue). (a) Lifetime of bulbous filopodia. (b) Total number of bulbous filopodia per terminal per hour. (c) Average number of bulbous filopodia per time instance. (d) Number of concurrently existing bulbous filopodia per axon terminal per time instance. (e, f) Representative snapshots of *lar* (e) and *trio* (f) revealing only transient bulbs. (g-n) Markov state model simulation for *lar* (g-j) and *trio* (k-n) for the numbers of filopodia (g, k), transient bulbs (h, l), stable bulbs (i, m) and synapses (j, n). In all cases control traces from Fig. 3 are shown in yellow. Black dotted lines: mean number of bulbs; solid red line: median number of bulbs; dark grey denotes the interquartile range (50% of the data) and light grey the 95% confidence range. (o-r) Measurement of BrpD3 punctae in mutant axon terminals. (o’-r’) BrpD3 single channel. Scale bar: 2 μm. (o-q) Measurement of BrpD3 punctae in *trio* and control axon terminals. (o’-p’) BrpD3 single channel. (q) Quantification of BrpD3-marked synapse numbers relative to control at P90. n=87 and 61, p= 0.67 (r) Schematic summary of protein functions during synapse formation.

Analysis of the bulb distribution at any given time point over a one hour period at P60 separates *lar* and *trio* further from *syd-1* and *liprin-α: lar* mutant terminals form no bulbs at most times points, supporting a defect in bulb initiation (**Fig. 5d**). In contrast, *trio* exhibits a wild type distribution suggesting that, in contrast to the other three mutants, one bulbous tip at any time point can still be stabilized (**Fig. 5d**). This is corroborated by live imaging (**Suppl. Movie 6**). The increased average bulb destabilization in *trio* is a function of many more bulbs forming, while only one appears stabilized at almost every time point. Taken together, our analyses of bulbous filopodia in the four mutants suggest that *lar* is defective in bulb initiation (**Fig. 5e**), *lar, syd-1* and *liprin-α,* fail to stabilize bulbs, and *trio* only exhibited destabilization of supernumerary bulbs (**Fig. 5f**).

We next used our complete time course simulation from filopodial dynamics to synapse formation using the measured P60 data for *lar* and *trio.* As shown in Figure 5g, simulated *lar* mutant terminals form normal numbers of filopodia. In contrast to *syd-1* and *liprin-α,* very few transient or stable bulbs form in *lar* mutants (**Fig. 5h,i**). Consequently, the *lar* simulation produces only very few synapses (**Fig. 5j**). In contrast, trio exhibits continuously elevated levels of filopodia, and increased number of transient bulbs (similar to *syd-1* and *liprin-α),* but close to wild type levels of stabilized bulbs (**Fig. 5m**), which lead to close to wild type levels of synapses (**Fig. 5n**). Correspondingly, BrpD3-labeling revealed normal numbers of synapses in *trio* (**Fig. 5o-p**). We could not reliably measure BrpD3-positive synapses in *lar,* because most axon are retracted by P70 and none retained a normal morphology.

In summary, our data reveal normal axon targeting and filopodial dynamics until P40 for all mutants, except for an increased filopodial ‘birth’ rate in *trio. lar* exhibits defective bulb initiation and *lar, syd-1* and *liprin-α* all fail to stabilize individual bulbs, leading to mostly transient and generally destabilized bulbs, many time points without bulbs, and significant reductions in synapse formation. In contrast, in *trio* axon terminals bulb destabilization almost exclusively occurs for supernumerary bulbs. This observation fits with a requirement of *trio* for negative feedback, but not bulb stabilization *per se.* In this scenario, a single bulb at any given time point still stabilizes, but fails to suppress other bulbs, resulting in supernumerary transient bulbs. Hence, all four mutants fit distinct roles of the ‘winner-takes-all’ mechanism and thus support the serial synapse formation model (**Fig. 5r**).

### A computational model predicts axon retractions in *lar, syd-1, and liprin-α,* but not in *trio*

We have so far performed all analyses on normally targeted axon terminals that remained stable in the correct target layers, thereby isolating defects in filopodial dynamics and synapse formation independent of axon retraction. Defects in bulbous filopodia stabilization for *lar, syd-1* and *liprin-α* mutant axons occurred at the correct target layer and thus prior to the possible later retractions of these axons. We therefore set out to test the idea that defective synapse formation might contribute to axonal destabilization of the initially correctly targeted axons. We have previously provided correlative evidence that a reduction in the dynamics of transient filopodia is associated with increased early retractions in *cadN* mutants^5^ In contrast to *cadN,* none of the mutants analyzed here exhibited altered filopodial dynamics or retractions prior to P40, when synaptic partner identification is likely to start. The first stable bulbous tips can be observed around P45 and synapse formation increases thereafter (**Fig. 6a**). Similar to filopodial adhesion, synapses may contribute to the stabilization of axon terminals. We therefore hypothesized that axon terminal stabilization may be a function of both a decreasing number of filopodia and increasing numbers of synapses (**Fig. 6a**).

**Figure 6:**
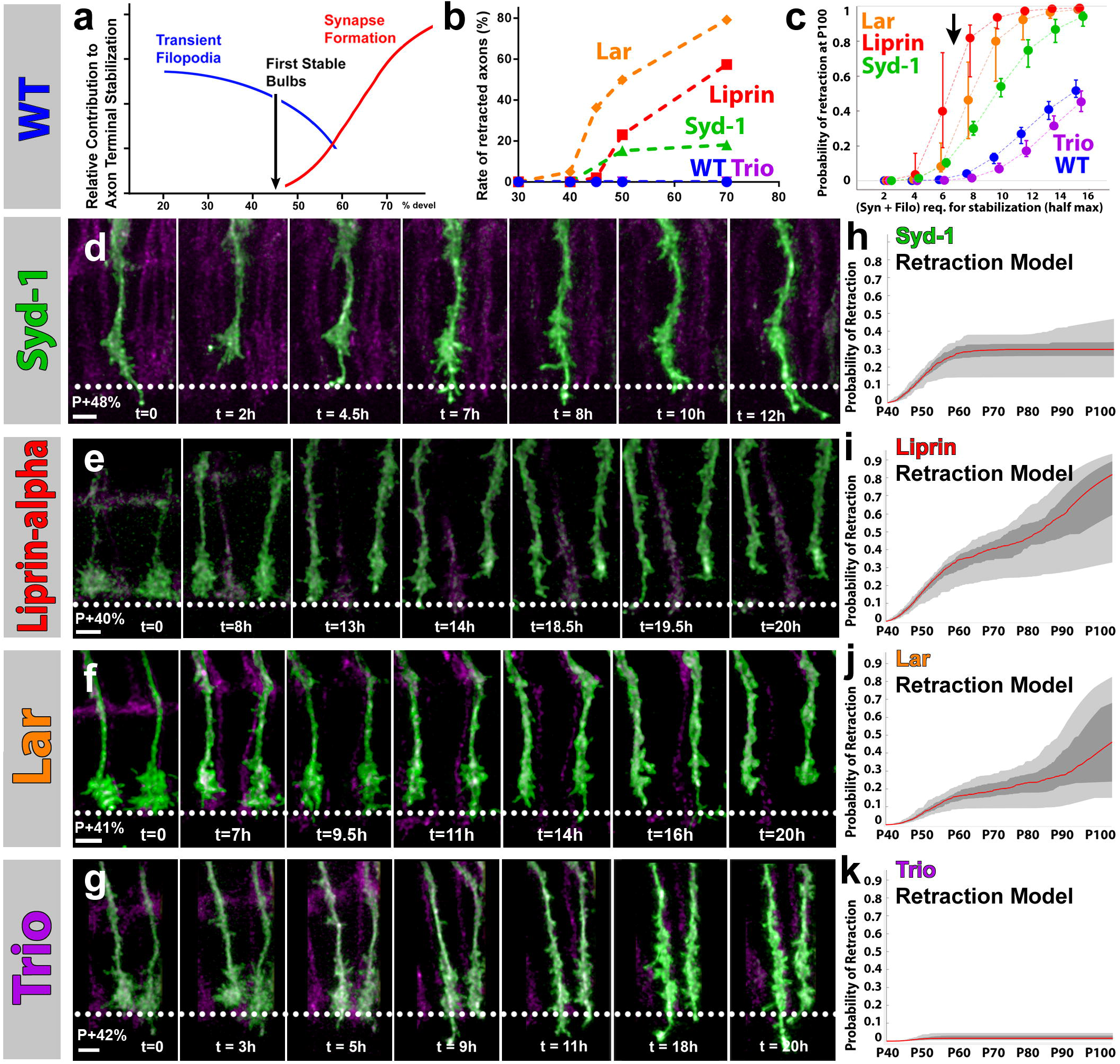
A computational model predicts axon retractions in *lar, syd-1, and liprin-α,* but not in *trio*. (a) Schematic of timeline during synapse formation, including continuous decline of transient filopodia, the first appearance of bulbs and the continuous increase in synapse numbers. (b) Measured R7 axon retraction rates. (c) Probability of R7 axon terminal retractions at P100 based on computational modeling of stabilization through a combination of transient filopodia and synapses. (d-g) Representative time-lapse snapshots from long-time live imaging of R7 axon stabilization and retraction in the four mutants. Dashed lines mark the wild-type R7 target layer (M6). Scale bars: 3 μm (h-k) Computational modeling of predicted probabilistic axon retractions between P40-P100 for all four mutants (comp. to measured data in panel (b).

First, we measured the retraction rates between P40-P70, revealing distinct properties for each of the four mutants (**Fig. 6b**). *lar* and *liprin-α* exhibit similar retraction rates with a 5- hour delay for *liprin-α* after *lar.* The dynamics of these retractions appeared similar in longterm live imaging of axon behaviors (**Fig. 6e-f, Suppl. Movie 7**). In both cases, individual terminals probabilistically collapse to a smooth structure within 2 hours and are not recognizably different just one hour prior. A filopodial protrusion often remained for several hours and the terminals retained the remarkable ability to re-extend to their correct target layer, but could not stabilize there. In contrast, apparent retraction of *syd-1* mutant axons plateau after P50 (**Fig. 6b**); *syd-1* axons initiated retractions very similar to *liprin-α* and *lar,* but exhibited many more re-extensions back to M6 and even beyond (**Fig. 6d, Suppl. Movie 7**). This behavior contributes to the appearance of less retracted *syd-1* axons after P50 (**Fig. 6b**). *trio* mutant axons exhibited increased filopodial extensions that are somewhat similar to *syd-1,* explaining the earlier observations of an additional overextension phenotype in fixed preparations of both *trio* and *syd-1* mutants^21^. We did not observe any retractions of *trio* mutant axons. However, careful analysis of *trio* mutant strains with the same genotypes as those used by Holbrook et al., 2012 revealed rare, misplaced R7 axons only in the original stocks from that study, but not in different genetic backgrounds. Hence, *trio* may have a mildly increased probability to retract that is only apparent depending on the genetic background.

To test whether these different retraction dynamics could in theory be the result of temporal changes in filopodial dynamics and synapse formation observed in the mutants, we performed data-driven quantitative retraction simulations. We modeled retraction probabilities as a function of the number of filopodia and synapses. If only filopodia stabilize axon terminals, but not synapses, the model predicts similar retraction rates for wild type and all mutants based on the measured filopodia numbers (**Suppl. Fig. 6**). In contrast, if synapses contribute to axon stabilization, the different synapse formation rates of the four mutants differentially affect retractions. The potential minimal number of filopodia or synapses required to stabilize an axon is unknown. We therefore tested the probability as a function of an equally weighted sum of filopodia and synapses based on the measured filopodia and simulated synapse formation data for all four mutants. If only very few filopodia or synapses are required to retain the axon, none of the mutants should exhibit axon retractions before P100 (**Fig. 6c**). If we increase the ‘minimal stabilization’ number, i.e. the number of filopodia plus synapses required to retain the axon, WT and all mutants exhibit an increasing probability to retract prior to P1 00. In this analysis, WT and *trio* exhibit the same low probability to retract only if high numbers of filopodia and synapses are required for stabilization (**Fig. 6c**). In contrast, *lar, liprin-α* and *syd-1* all exhibit significantly increased probabilities to retract. Notably, the regime where only *lar, liprin-α* and *syd-1* exhibit retractions is robust over a wide range of the ‘minimal stabilization factor’ (**Fig. 6c**). As shown in Figures 6h-k simulation of the retraction dynamics of all mutants for the ‘minimal stabilization’ number marked by an arrow in Figure 6c. Remarkably, all four mutants exhibit retraction kinetics that closely resemble the observed retractions. In particular, *liprin-α* exhibits only slowly decreasing retraction rates, while *syd-1* appears significantly more dampened after P60 (comp. Fig. 6b and 6i, j). The least good match is *lar.* the data show significantly earlier retractions with higher rate than the model. This suggests that retractions in *lar* are not sufficiently explained as a function of loss of synapses, but may occur earlier already due to an additional adhesion role, as previously suggested^31,32^. In sum, our combined live dynamics measurements and data-driven modeling suggest that the serial synapse formation model is sufficient to predict the number and distribution of synapses and their role in stabilizing axon terminals in wild type and *liprin-α, syd-1* and *trio.* Our data further suggest that Lar plays a role in the same process as Liprin-α and Syd-1, but early retractions may be caused by an additional, earlier function.

## Discussion

In this study we characterized the role of filopodial dynamics during synapse formation using the *Drosophila* R7 photoreceptor terminal as a model. We identified a rare type of filopodium with enlarged, bulbous tips, that only occur during synapse formation. Only a single bulbous filopodium per axon terminals is stable at a given time point. The early synaptic assembly factors Syd-1 and Liprin-α similarly distribute in only one filopodium at a time. Loss of these factors affects the stabilization of bulbous filopodia, while the vast majority of filopodial dynamics throughout development are unaffected. As a consequence, development is normal until the time of synapse formation. Thereafter, these mutants fail to stabilize bulbous filopodia, leading to a failure to form synapses and the probabilistic destabilization of axon terminals. In addition, measurements and modeling of *lar* and *trio* mutants indicate a role in bulb initiation (*lar*) and the implementation of suppressive feedback on other bulbs following single bulb stabilization (*trio*). These findings suggest a serial synapse formation model based on competitive distribution of synaptic building material between specialized filopodia (**Fig. 7**).

**Figure 7:**
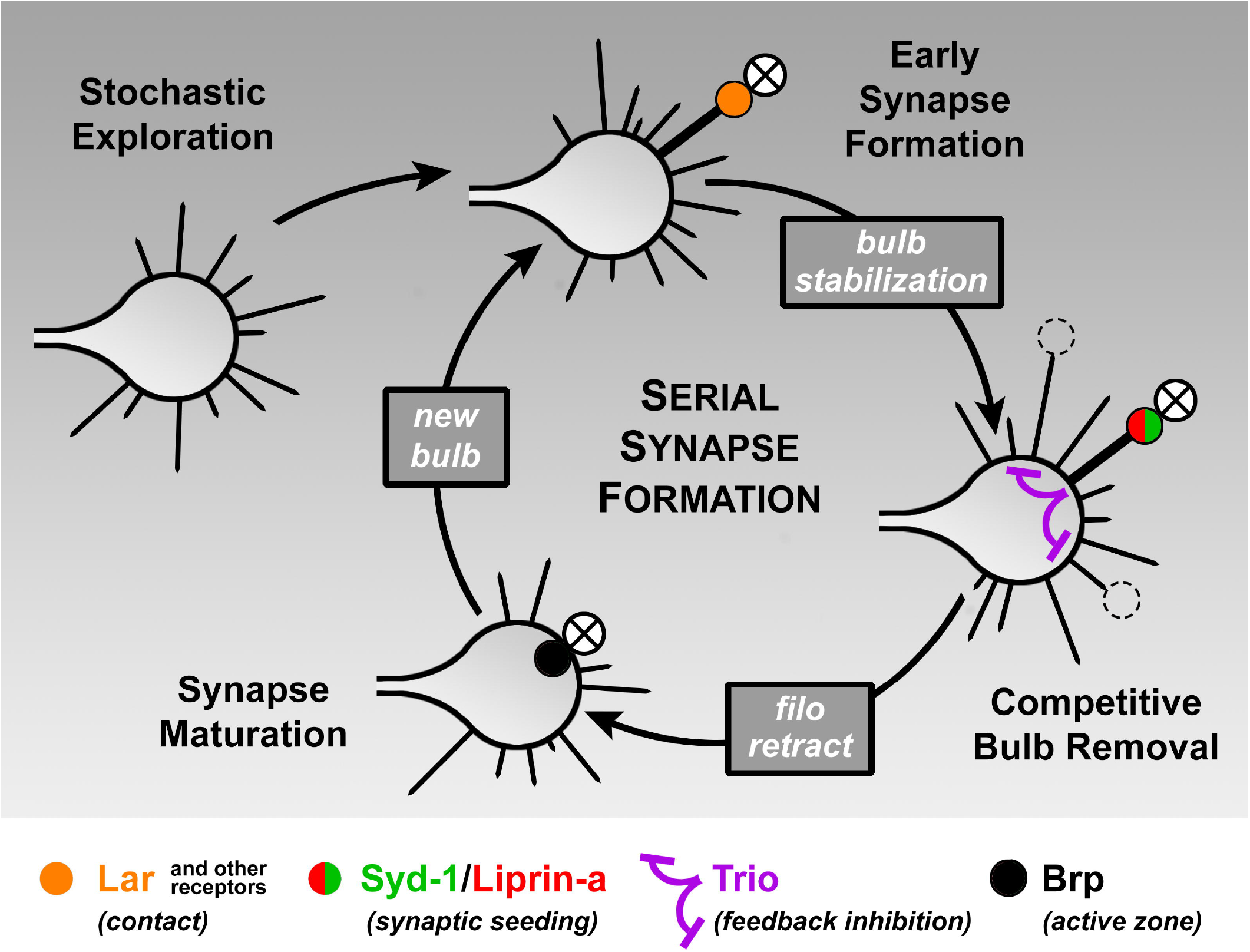
Serial Synapse Formation Model. The measured live dynamics and computational modeling in this paper suggest the following model: (1) stochastic filopodial exploration leads to synaptic capture via a cell surface receptor, e.g. lar; (2) early synaptic seeding factors (Syd-1 and Liprin-α) are recruited to the captured filopodium in an enlarged bulb; (3) secondary simultaneously forming bulbs are destabilized via the function of the RhoGEF Trio, thereby ensuring one synaptogenic filopodium at any given time; recruitment of the active zone protein Brp and synapse maturation occur after filopodial retraction back in the main axon terminal, allowing a new cycle of bulb formation an stabilization.

### Serial Synapse Formation through filopodial competition for synaptic seeding factors

Our 4D tracking and quantitative analysis of filopodial dynamics in wild type and synapse formation mutants led to the characterization of a rare filopodium that formed bulbous tips. Our data link bulbous filopodia to synapse formation based on three findings: (1) In wild type, these are the only filopodia that only occur during the time window of synapse formation and do not exhibit stochastic dynamics; wild type R7 photoreceptors stabilize one bulbous filopodium at a time. (2) The synaptic seeding factors Liprin-α and Syd-1 non-randomly localize to one bulbous filopodium at a time. (3) Loss of *liprin-α* or *syd-1* selectively affects the stabilization of bulbous filopodia, but no other filopodial dynamics prior to synapse formation. In addition, loss of the upstream receptor *lar* similarly selectively affects bulbous filopodia, but, in addition to bulb destabilization, also strongly affects bulb initiation. Together, these findings support a model whereby stochastic filopodial exploration leads to bulb formation and stabilization one at a time, which in turn leads to the formation of one synapse at a time. This model effectively controls synapse numbers within the available developmental time window.

The key mechanism of this model is inhibitory feedback of bulb formation. In contrast to all other filopodia, the dynamics of bulbous filopodia are not independent. We hypothesize that competitive bulb stabilization depends on synaptic seeding factors, because Syd-1 and Liprin-α accumulate only in one bulb at a time and their loss leads to bulb destabilization. How are synaptic seeding factors competitively and non-randomly distributed between filopodia? Our live imaging data suggest larger aggregates of Liprin-α or Syd-1 can traffic in and out of filopodia, while the mature active zone protein Brp is prevented from entering filopodia. Overexpressed GFP-Syd-1 and Liprin-α-GFP accumulate in the axon terminal trunk and do not lead to more than one filopodium containing larger amounts of either protein. These observations indicate that trafficking into filopodia is restricted. Morphologically, filopodia are very thin structures that may not provide much space for freely diffusing cytosolic proteins, aggregates or organelles. On the other hand, the bulbous tip provides a much larger volume that may be required for sufficient amounts of synaptic seeding factors and other building material to initiate synapse formation.

Since Syd-1 and Liprin-α are not required for bulb initiation, we speculate that filopodial contact with a synaptic partner may initiate the bulb and precede active zone assembly through seeding factors in the presynaptic cell. Our data suggest that Lar is a good candidate for a presynaptic receptor with such a role, but is unlikely to be the sole upstream receptor. Neurexin^46^ and PTP69D^47,48^, for example, are other known candidates. In the absence of an upstream receptor that keeps seeding factors at the membrane or the seeding factors themselves, synapse assembly fails and bulbs destabilize. The significantly increased frequency of new bulb generation in mutants for the seeding factors suggests a compensatory mechanism. Hence, both the non-random competitive stabilization of single bulbs in the presence of seeding factors, as well as the immediate new bulb generation following loss of bulbs in the absence of seeding factors imply terminal-wide communication of the presence of stable bulbs. This is reminiscent of other competitive processes that shape neuronal morphology, e.g. the restricting role of building material in the competitive development of dendritic branches in a motorneuron^49^. Our mutant analyses suggest that stable bulbs communicate negative feedback to other bulbs via the function of the RhoGEF *Trio.* While the exact mechanism is unclear, it is tempting to speculate about a role of actin-dependent signaling downstream of synaptic seeding.

Note that the presence of a stable bulb does not prevent the initiation of a new bulb, it only prevents seeding factor-dependent stabilization of the competitor. Hence, the competition is independent of the initiation of bulbs and thus likely synaptic contacts. This idea is also supported by the observation that the process of serial synapse formation is robust to an increase in filopodia dynamics and bulb formation, as is observed in the *trio* mutant, because the stabilization of only one bulb at a time ensures quantitatively robust synapse formation.

### Cause and Effect: The Challenge to Identify Primary Defects in Circuit Assembly

Mutations in the proposed pathway components Lar, Liprin-α, Syd1 and Trio have been independently characterized for their roles in active zone assembly (mostly at the larval neuromuscular junction) and axon targeting, in large part in the visual system^20,21,27,31,32^. It is likely that all four genes exert more than one function in different contexts. Especially for Lar, independent context-dependent function have been characterized based on different downstream adaptors^31^. In our study, we asked to what extent a primary role of these genes in synapse formation in *Drosophila* photoreceptors could explain previously observed phenotypes. All filopodial defects measured here occur independent and prior to possible retraction events. Our combined live imaging and computational modeling approach suggests that defects in the *syd-1* and *liprin-α* mutant are consistent with a primary defect in bulb stabilization and synapse formation. These defects in bulb dynamics and synapse formation may in turn lead to axon destabilization or represent independent functions; *lar* may have an additional earlier adhesion function and *trio* does not play a critical role in the formation of the correct number of synapses, while its effect on general filopodial dynamics may sensitize mutant axons to other changes.

We base our conclusion that Lar, Liprin-α and Syd-1 have a primary function in synapse formation on three pieces of evidence: (1) All mutants initially target correctly and exhibit normal filopodial dynamics prior to synapse formation; (2) All mutants start retracting only when synaptic contacts initiate, in the order and severity from the receptor to the downstream elements; *lar* and *liprin-α* track each other with similar retraction rates; (3) All three mutants cause the loss of competitive bulb stabilization. Taken together, these observations support a direct role in synapse formation following bulb stabilization. However, we cannot exclude other molecular functions for any of these proteins. For example, in both *C. elegans* and *Drosophila* Lar has been shown to function independently in axon guidance and synapse formation^31,50^

Interestingly, we found that *syd-1^ΔRhoGAP^* mutants have normal terminal morphology and only a mild decrease in the number of BrpD3-puncta. This is consistent with recent findings that a RhoGAP-deficient Syd-1 fragment is sufficient rescue early active zone seeding events at the NMJ but not the recruitment of Brp as the active zones mature^22^. However, since homozygous *syd-1^ΔRhoGAP^* flies are viable and fertile without obvious connectivity defects, synapse numbers are apparently sufficient for axon terminal stabilization.

Finally, our observations suggest that the primary functions in filopodial dynamics and synapse formation are sufficient to cause axon retractions. We have previously shown that loss of N-Cadherin leads to probabilistic retractions and re-extensions of R7 terminals long before synapse formation. The phenotypes observed here for lar, *liprin-α* and *syd-1* are reminiscent, but only occur at or after the time of synaptic partner identification. We tried to combine stabilization through filopodial adhesion and synapse formation giving equal weights to both (**Fig. 6a**); while filopodia continuously decrease, synapses continuously increase, thereby allowing a take-over of the stabilization function. The modelling fits wild type, *liprin-α, syd-1* and *trio* remarkably well. On the other hand, retractions in the *lar* mutant are qualitatively predicted, but the model fails to explain retractions quantitatively. A partial explanation may be that we parameterized our model only based on the *lar* mutant axon terminals that are still unretracted at P60. These are only 30% of terminals by that time, and we have effectively selected for terminals with dynamics that prevented retractions thus far. It is likely that earlier retractions are caused by defects in filopodial adhesion or synaptic contacts.

## Methods

### Molecular Biology

To build the UAS-BrpD3-mKate2 construct, EGFP sequence was removed from the pTW BrpD3-GFP plasmid (gift from S. Sigrist) using Xba1 and Age1 sites. mKate2 sequence was amplified from the pmKate2-C plasmid (Evrogen) using the following forward and reverse primers (respectively): gggTCTAGACggtggaggaggtATGGTGAGCGAGCTGATTAA and cccACCGGTTTATCTGTGCCCCAGTTTGCTAG. The products were digested with Xba1 and Age1 and ligated into the above-mentioned pTW BrpD3 plasmid. Injections were done by Rainbow Transgenics (USA) for P-element insertion and candidate lines were isolated and tested according to standard procedures.

*syd-1*^dRhoGAP^ allele was generated by Well Genetics (Taiwan) using CRISPR/Cas9 Scarless (DsRed) system (**Suppl. Fig. 4a**). 2 gRNAs were used against the following target sites (PAM): CGGGAGTCTAAGAATGCTCC[CGG]; AGATACTTAAGCACCGCGAT[CGG]. Upon PBac-mediated excision, a specific and complete deletion of the RhoGAP domain was achieved with only a TTAA motif left embedded in the exogenous sequence GTTAAA (Fig. SXB). Insertion and excisions were verified by genomic PCR and sequencing. Full design details and sequencing results are available upon request.

### Genetics

All experiments were performed with *Drosophila* pupae collected at P+0% (white pupae) and aged in 25°C unless otherwise specified. The following *Drosophila* genotypes were used. For wild-type membrane and synapse imaging: (GMR-FLP/+; GMR-Gal4/ GMR-myr-tdTomato; FRT80B, UAS-CD4-tdGFP/ FRT80B, tub-Gal80) and (GMR-FLP/+; FRT42D, GMR-Gal80/ FRT42D; GMR-Gal4, UAS-CD4-tdTomato/ UAS-BrpD3-GFP). Membrane imaging with mutants: (GMR-FLP/+; FRT40A, tub-Gal80/ FRT40A, *liprin-α^E^* (or dlar^2127^); GMR-Gal4, UAS-CD4-tdGFP, GMR-myr-tdTomato/+), (GMR-FLP/+; GMR-Gal4, UAS-CD4-tdGFP/ GMR-myr-tdTomato; FRT82B, tub-Gal80/ FRT82B, syd-1^w46^ (or syd-1^dRhoGAP^)), (GMR-FLP/+; GMR-Gal4, UAS-CD4-tdGFP/ GMR-myr-tdTomato; FRT2A, tub-Gal80/ FRT2A, *trio*^3^). Synaptic imaging with mutants and corresponding controls: (GMR-FLP/+; FRT40A, tub-Gal80/ FRT40A, *liprin-α*^E^ (or dlar^2127^ or FRT40A only); GMR-Gal4, UAS-CD4-tdTomato/ UAS-BrpD3-GFP), (GMR-FLP/+; GMR-Gal4, UAS-CD4-tdGFP/ UAS-BrpD3-mKate2; FRT82B, tub-Gal80/ FRT82B, syd-1^w46^ (or syd-1^dRhoGAP^ or FRT82B only)), (GMR-FLP/+; GMR-Gal4, UAS-CD4-tdGFP/ UAS-BrpD3-mKate2; FRT2A, tub-Gal80/ FRT2A, *trio*^3^ (or FRT2A only)). For imaging of early synaptic markers: (GMR-FLP/+; GMR-Gal4; FRT80B/ UAS-Liprinα-GFP (or UAS-GFP-Syd1), UAS-CD4-tdTomato/ FRT80B, tub-Gal80). For imaging with Brp RNAi: (GMR-FLP/+; FRT42D, GMR-Gal80/ FRT42D; GMR-Gal4, UAS-CD4-tdGFP, GMR-myr-tdTomato/ UAS-Brp-RNAi^B3^, UAS-Brp-RNAi^C8^). For ERG recordings: (; GMR-Gal4/ FRT42D;) and (; GMR-Gal4/ FRT42D; UAS-Brp-RNAi^B3^, UAS-Brp-RNAi^C8^/ +). Sources for all transgenic flies are listed in **Supplementary Table 2**.

### Histology and Fixed Imaging

Eye-brain complexes were dissected in PBS, fixed in 3.7% paraformaldehyde (PFA) in PBS for 40 minutes, washed in PBST (0.4% Triton-X) and mounted in Vectashield (Vector Laboratories, CA). Images were collected using a Leica TCS SP8-X white light laser confocal microscope with a 63X glycerol objective (NA=1.3).

### Brain Culture and Live Imaging

*Ex vivo* eye-brain complexes were prepared as described before^5^. For filopodial imaging, brains were dissected at P+40% and 1 μg/ml 20-Hydroxyecdysone was included in the culture media. For synaptic imaging, brains were dissected at P+50% and no ecdysone was included.

Live imaging was performed using a Leica SP8 MP microscope with a 40X IRAPO water objective (NA=1.1) with a Chameleon Ti:Sapphire laser and Optical Parametric Oscillator (Coherent). We used a single excitation laser at 950 nm for two-color GFP/Tomato imaging. For GFP/mKate2 imaging lasers were set to 890 nm (pump) and 1150 nm (OPO).

### Electroretinogram (ERG) Recordings

1-5 day-old adult flies were reversibly glued on slides using nontoxic school glue. Flies were exposed to 1s pulses of light stimulus provided by computer-controlled white light-emitting diode system (MC1500; Schott) as previously reported^51^. ERGs were recorded using Clampex (Axon Instruments) and measured using Clampfit (Axon Instruments).

### Data Analysis

All live imaging data as well as all data involving synaptic markers were deconvolved (10 iterations with the theoretical PSF) using Microvolution Fiji Extension. Imaging data were analyzed and presented with Imaris (Bitplane). For synaptic counts, Spot objects were created from the BrpD3 channel and Surfaces were generated from the CD4 channel using identical parameters between experimental conditions and the corresponding control. Spots were then filtered for their localization on the positive clones by the Imaris 9 MATLAB extension ‘XTSpotsCloseToSurface’.

Further analysis regarding the quantified data and generation of corresponding graphs were done using Prism 7 (GraphPad). Where needed statistical differences were calculated with unpaired, parametric t-tests.

### Filopodia Tracing and Tracking

We developed an extension to the Amira Filament Editor^35^. for tracing and tracking of individual filopodia in 4D datasets. Growth cones are represented by an annotated skeleton tree, in which each branch corresponds to a filopodium (**Suppl. Fig. 1b**). This tree is traced for each time step and matched to the tree in the previous time step in a semi-automatic process.

First, the user interactively marks the growth cone (GC) centers in the first time step. The GC centers are automatically detected in the remaining time steps using template matching^52^ Then, the GCs are processed one at a time. To this end, the images are cropped such that they contain only the current GC. The user interactively specifies the filopodia tips in the first time step. The filopodia are traced automatically from the tip to the GC center using an intensity-weighted Dijkstra shortest path algorithm based on^53^. The onset of a filopodium is determined by identifying the point on the path where the 2D intensity profile orthogonal to the tracing changes from Gaussian (for the filopodium) to non-Gaussian (inside the GC body). The user visually verifies the tracing and, if necessary, interactively corrects it using dedicated tools provided by the Filament Editor. After tracing all filopodia in the first time step, they are automatically propagated to the next time step by template matching of tips and onsets, and tracing paths from tip to center through the onset. Propagated filopodia obtain the same track ID as the original. After each time step the user verifies the generated tracings, and adds newly emerging filopodia. This process is continued until all time steps have been processed (**Suppl. Fig. 1a**).

Statistical quantities including length, angle, extension/retraction events, and lifetime are extracted from the filopodia geometry and stored in spreadsheets.

### Mathematical Modeling

Stochastic filopodial dynamics were modelled by a Markov jump (Poisson-) process formalism as outlined in the Supplementary Note. Briefly, we developed a data-driven minimal model capturing the dynamics of filopodia, bulbous tips and synapse formation. We first identified the systems variables and subsequently estimated model parameters, also testing higher complexity models whenever the best fit of a simpler model could not sufficiently explain the data. All data used for model inference and parameterization, as well as the parameter inference procedure are exemplified in the Supplementary Note. All codes were written in MATLAB 2018a (Mathworks, Nattick). Parameter inference was performed using the MATLAB 2018a function ‘fminsearch’ and simulations were performed using the stochastic simulation algorithm. For detailed descriptions of all procedures see the Suppl. Notes on Mathematical Modeling.

## Acknowledgements

We would like to thank all members of the Hiesinger, Wernet and Hassan labs for their support and helpful discussions. We thank Stephan Sigrist and Claude Desplan for critical reading of the manuscript. We further thank Thomas Clandinin, Larry Zipursky and Stephan Sigrist for reagents. This work was supported by the NIH (RO1EY018884, RO1EY023333) and the German Research Foundation (DFG) (SFB 958, SFB186) and FU Berlin. Max von Kleist acknowledges financial support from the BMBF grant number 031A307 and Marian Moldenhauer, Martin Weiser and Max von Kleist acknowledge support from MATHEON and the Einstein Center for Mathematics Berlin, provided through the Einstein Stiftung Berlin.

