## Supplemental Note, Figures and Tables for "Serial synapse formation through filopodial competition for synaptic seeding factors"


| | |
|---|---|
| Supplementary Note on Mathematical Modeling | 1-18 |
| Supplementary Figures 1-6 and Legends | 19-24 |
| Supplementary Table 1 | 25 |
| Supplementary Table 2 | 26 |

### Supplementary Note: Mathematical Modelling.

In this supplement we describe the development of a minimal and sufficient mathematical model describing stochastic filopodial dynamics and synapse formation. Specifically, the model is data-driven and based on measurements of the rates of all observed filopodia and the quantitative emergence of synapses between Drosophila pupal stages 40% and 100% of development. We first built a reference model for wild type and then adapted it to the four knock-out mutants syd-1, liprin-alpha, lar and trio in a data-driven fashion.

#### SN.1 Markov jump model.

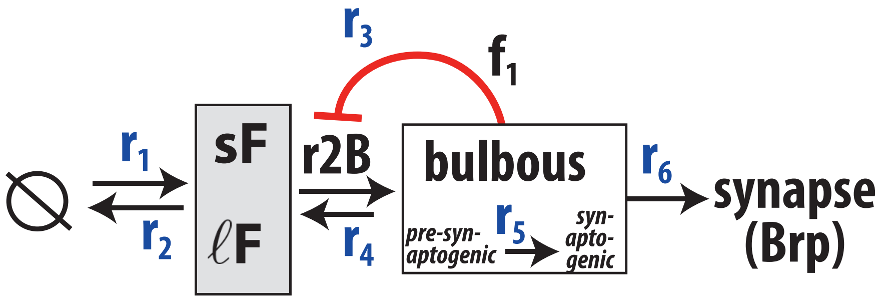


Figure SN.1: Schematic of the stochastic model for the filopodial dynamics (including short-lived $sF$ and long-lived $\ell F$ filopodia), bulbous $B$ and synapse $S$ formation. Reactions $r_1$ and $r_2$ denote the 'birth' and retraction of filopodia. Formation of a bulbous tip $B$ and the retraction of a bulbous tip filopodium are denoted by $r_3$ and $r_4$ respectively. The rate at which a bulbous tip becomes synaptogenic (stabilized) is denoted by $r_5$ and $r_6$ is the rate at which synaptogenic bulbous filopodia become synapses $S$.

The observed stochastic dynamics of filopodia appearance and retraction (Fig.1 E, main text) prompted us to use a Poisson process formalism. Foremost, our intention was to identify the state variables of this system. Besides synapses ($S$), which denote the endpoint of the modelling pipeline, bulbous tips were directly identified in the time-lapse data based on their altered morphology. Analysis of the bulbous life time data in the mutants identified two populations: short-lived, unstable bulbous tips ($sB$) that appeared and disappeared within the 60 minutes imaging interval vs. stable bulbous tips that persisted once appeared (Fig. SN.4). We called the latter type 'synaptogenic bulbous tips' ($synB$). Interestingly, however, the unstable bulbous population ($sB$) was almost absent in the wild type (more below). Moreover, we identified two types of filopodia, which are distinguished by their lifetime and which will henceforth be denoted short-lived- ($sF$) and long-lived ($\ell F$) filopodia. The final model is depicted in Fig. SN.1, and the reaction stoichiometries are determined by the following reaction scheme:

$$R_{1,sF} : \emptyset \longrightarrow sF \quad , \quad R_{2,sF} : sF \longrightarrow \emptyset \quad , \quad R_{1,\ell F} : \emptyset \longrightarrow \ell F \quad , \quad R_{2,\ell F} : \ell F \longrightarrow \emptyset \qquad \text{(SN.1)}$$

$$R_3 : F \to sB \quad , \quad R_4 : sB \to \emptyset \quad , \quad R_5 : sB \to synB \quad , \quad R_6 : synB \to S \qquad \text{(SN.2)}$$

Note that in $R_3$ we denote by $F$ any filopodium (short-lived and long-lived) and in $R_4$ we have ignored the flux back into the filopodia compartment $sF + \ell F$ as it insignificantly affects the number number of filopodia (small $B$, small rate $r_4$), as will become evident in this note.
Below, we will guide through the model building and parametrisation process.

##### SN.1.1 Model building

**Short-lived vs. long-lived filopodia.** Our aim was to develop a minimal model of filopodial dynamics and synapse development. Besides the state variables for synapse $S$ and bulbous tips $B$, which can be directly identified in the data, it is unclear whether a single- or multiple morphologically indistinguishable filopodia types may be present. 4D filopodia tracking allows to generate thousands of trajectories of individual filopodia during high-resolution recordings at P40 and P60 respectively (Supplementary Figure S1). From this trajectory data we can compute the statistics of individual filopodial life times (Fig. SN.2). We then asked whether a single filopodia compartment, or two distinct filopodial sub-populations, would reproduce this life time data. The figures below (Fig. SN.2) show the respective fits if we considered one- vs. two filopodia compartments, assuming an exponential life time distribution respectively. As can be seen, the data strongly supports the existence of two filopodial compartments, based on the life time data ($AIC_{2cmp}$ = 260 (three parameters) versus $AIC_{1cmp}$ = 827 (one parameter)). The respective rate constants for retraction were $c_{2,sF}$ = 0.69 ($min^{-1}$) and $c_{2,\ell F}$ = 0.12 ($min^{-1}$) for short- and long-lived filopodia. The optimal cut-off to differentiate these two populations in the data based on their lifetimes was 8

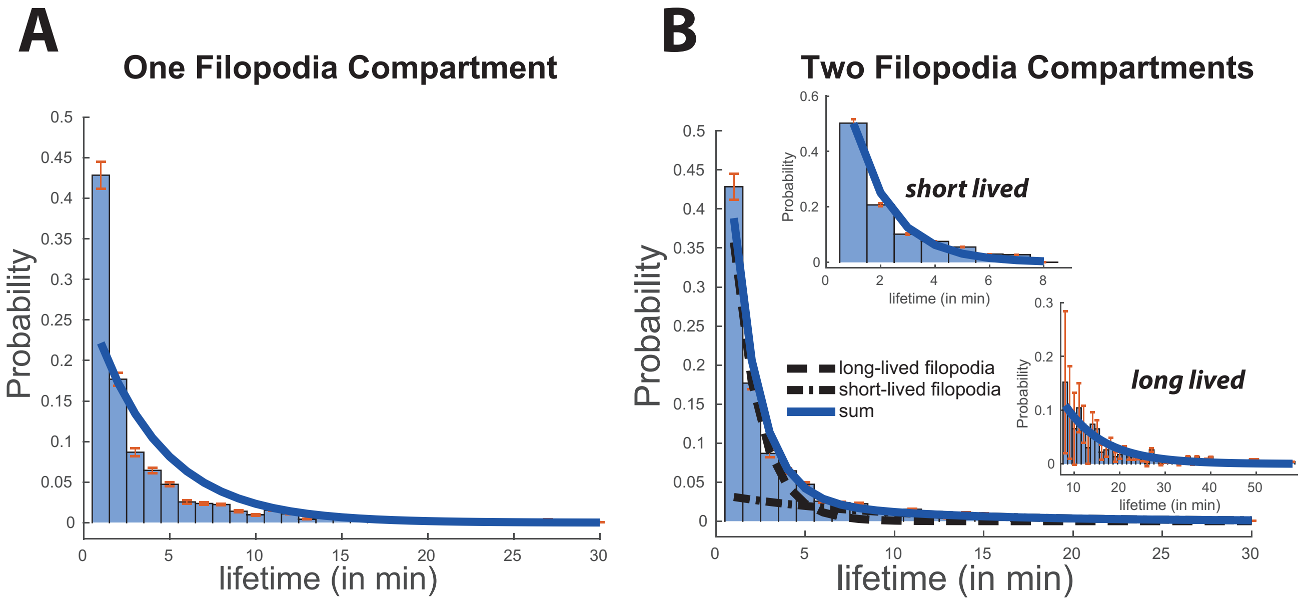


Figure SN.2: Filopodial life times and identification of two filopodia subpopulations. Blue bars indicate the empirical filopodia life time distribution from individual filopodia recordings at P40 and P60 respectively. **A.** The thick blue line indicates the best fit assuming a single filopodium population with an exponential lifetime. **B.** The thick blue line indicates the best fit assuming two filopodium sub-populations with distinct exponential lifetimes. The short-lived filopodia ($sF$) retract with rate constant 0.69 ($\text{min}^{-1}$), whereas the fast filopodia retract with rate 0.12 ($\text{min}^{-1}$).

minutes, Fig. SN.2B (inset).

**Transient vs. stable (synaptogenic) bulbous tip filopodia.** Analysis of the bulbous life time data *in the mutants* identified two populations: short-lived, unstable bulbous tips ($sB$) that appeared and disappeared within the 60 minutes imaging interval vs. stable bulbous tips that persisted once appeared (Fig. SN.4). We called the latter type 'synaptogenic bulbous tips' ($synB$), assuming that materials necessary to form synapses require time to accumulate in bulbous tips, also stabilising them. Interestingly, in the wild type almost all bulbous tips were stable in terms of prolonged life time. For the mutants, the life time distribution of the short-lived (transient) bulbous tips (left bars in Fig. SN.4) appeared exponentially distributed with a mean life time of 3.3, 9, 5.9 and 7.6 *min* for the lar, liprin-alpha, syd-1 and trio mutants. As mentioned, in the wild type we did observe transient bulbous tips and assumed that they have a mean half life of about 120 minutes (in consequence most bulbous tips in the wild type will eventually become synaptogenic).

#### SN.1.2 Model Parametrisation

**Retraction and generation of filopodia, $r_1, r_2$.** As outlined in the previous paragraph, we identified two filopodia populations with exponential life times respectively. The exponential life times indicate a first-order decay with the respective rate constants $c_{2,sF} = 0.69$ ($\text{min}^{-1}$) and $c_{2,\ell F} = 0.12$ ($\text{min}^{-1}$) as derived in the previous section.

Additionally, we observe in Fig.1F (main article) that the number of filopodia per time instance is Poisson distributed, e.g. $sF \sim \mathcal{P}(\lambda_{sF})$ and $\ell F \sim \mathcal{P}(\lambda_{\ell F})$, where $\lambda$ denotes the average number of filopodia per time instance. Given the first-order retraction of filopodia, the Poisson distribution can be explained by a zero-order input with rate $c_{1,sF}$ and $c_{1,\ell F}$ and $\lambda_{sF} = c_{1,sF}/c_{2,sF}$ and $\lambda_{\ell F} = c_{1,\ell F}/c_{2,\ell F}$ respectively. The latter is a well-established result, e.g. regarding the stationary distribution of a birth-death process [1].

Interestingly, while the life time data revealed no significant changes in $c_2$ using data from only P60 vs. the pooled sample, the average number of filopodia was significantly different and in fact decreased significantly over the the 20 hours window from P40 to P60, Fig.1F(main manuscript) and Table SN.5. This prompted us to introduce a time-dependent function $f_F(t)$ that down-regulates the generation of new filopodia at a slow time scale. The time-dependent function $f_F(t)$ was then fitted to normalized filopodia counts at P40–P100, as shown in Fig 3B (main text). In summary, the propensity functions for reactions $R_{1,sF}$, $R_{2,sF}$,$R_{1,\ell F}$, $R_{2,\ell F}$ are given as follows.

$$r_{1,sF}(t) = f_F(t) \cdot c_{1,sF} \qquad r_{2,sF}(sF) = sF \cdot c_{2,sF} \tag{SN.3}$$

$$r_{1,\ell F}(t) = f_F(t) \cdot c_{1,\ell F} \qquad r_{2,\ell F}(F) = \ell F \cdot c_{2,\ell F} \tag{SN.4}$$

where $f_F(t) = \max(0, \sum_{i=0}^{5} p_i \cdot t^i)$ is a fifth-order polynome with coefficients $p_5 = -2.97 \cdot 10^{-14}, p_4 = 3.31 \cdot 10^{-13}, p_3 = -1.29 \cdot 10^{-9}, p_2 = 2.06 \cdot 10^{-6}, p_1 = -1.45 \cdot 10^{-3}$ and $p_0 = 1$. Note, that $t$ denotes the time in (min) *after* P40 (e.g. $t_{P40} = 0$).

| WT | P40 | P60 | $P$-value |
|---|---|---|---|
| $sF$ | 5.6 | 3.4 | p < 0.001 |
| $\ell F$ | 10 | 4.9 | p < 0.001 |

Table SN.1: Average numbers of filopodia per time instance. $P$-values were computed using the bootstrap technique: To generate a bootstrap sample, we drew 60 imaging frames with replacement at P40 and P60 respectively. We then e.g. counted the number of bootstrap occurrences $\lambda_{sF,P40} \leq \lambda_{sF,P60}$, denoted $X_{\lambda_{sF,P40} \leq \lambda_{sF,P60}}$. The p-value for assessing $\mathcal{H}_0 : \lambda_{sF,P40} \leq \lambda_{sF,P60}$ vs. $\mathcal{H}_1 : \lambda_{sF,P40} > \lambda_{sF,P60}$ is then given by $P = X_{\lambda_{sF,P40} \leq \lambda_{sF,P60}}/B$, where $B = 5000$ is the number of bootstrap samples.

Consequently, we have $f_F(t_{P40}) = 1$ and we can determine the input rate constant directly from the average number of filopodia at P40, i.e. $c_{1,sF} = \lambda_{sF,P40} \cdot c_{2,sF}$ and $c_{1,\ell F} = \lambda_{\ell F,P40} \cdot c_{2,\ell F}$ respectively. Raw filopodia counts are depicted in Fig. SN.5–SN.6 at the end of this document.

**Bulbous dynamics, $r_3, r_4$ and $r_5$.** Foremost, we assumed that short-lived unstable bulbous tips retracted by first order kinetics (reaction $r_4$). Using this assumption, the rate constant of retraction is equal to the inverse of the expected lifetimes of bulbous tips, which are stated in Table SN.3 at the end of this document.

We then wanted to investigate whether the bulbous tip number distributions in Fig. SN.8 and Fig. SN.10 can be explained by simple input-output relations or whether regulatory/feedback mechanisms are involved.
The number distribution of short-lived bulbous tips $sB$ and synaptogenic (stabilized) bulbous tips $synB$ is given by:

$$* \underset{r_4}{\overset{r_3}{\rightleftharpoons}} sB \underset{r_5}{\longrightarrow} synB \underset{r_6}{\longrightarrow} * \tag{SN.5}$$

**Model I: No feedback:** In the absence of any regulatory mechanisms (feedbacks), all reaction rates are of first order, e.g. $r_4 = sB \cdot c_4$, $r_5 = sB \cdot c_5$, and $r_6 = synB \cdot c_6$. The *net* influx $r_3$ at $t$ =P60 is

$$r_3(t) = c_3 \cdot (sF(t) + \ell F(t)) \cdot f_{FB}(t) \tag{SN.6}$$

where we assume that $sF(t), \ell F(t)$ and $f_{FB}(t)$ are approximately constant over the time scale of interest. Parameters $c_4$ can be approximated by (the inverse of) the bulbous tip life times presented in Table SN.3 and $c_6 \approx 1/120$ (min$^{-1}$) can be approximated from the maximum slope of synapse generation presented in Figure 3H (main manuscript), assuming the serial synapse formation model (at most one synapse can be formed at a time). The two parameters $c_5$ and $r_3(t)$ remain to be estimated for $t = P60$.
To perform this task we set up a generator matrix $G$ that has entries (transition rates)

$$G([i, j], [i-1, j]) = i \cdot c_4, \qquad G([i, j], [i, j-1]) = j \cdot c_6 \tag{SN.7}$$

$$G([i, j], [i+1, j]) = r_3(t), \qquad G([i, j], [i, j+1]) = i \cdot c_5 \tag{SN.8}$$

and diagonal elements such that the row sum equals 0. In the notation above, the tupel $[i, j]$ denotes the state where $i$ short-lived bulbous tips $sB$ and $j$ synaptogenic bulbous tips $synB$ are present. The generator above has a reflecting boundary at sufficiently large $N$ (maximum number of bulbous tips). The stationary distribution of this model is derived by solving the eigenvalue problem

$$G^T \cdot v = v \cdot \lambda$$

and finding the eigenvector corresponding to eigenvalue $\lambda_0 = 0$. From this stationary distribution, we compute the marginal densities of $sB$ and $synB$ (e.g. summing over all states where $i = 0, 1, ...$ for $sB$) and fit them to the experimentally derived frequencies in Fig. SN.8 and Fig. SN.10 by minimizing the Kullback-Leibler divergence between the experimental and model-predicted distributions. The resulting best fit for the wild type is shown in Fig SN.3 (dashed lines).
Lastly, parameter $c_3$ is derived by calculating

$$c_3 = \frac{r_3(t)}{(sF(t) + \ell F(t)) \cdot f_{FB}(t)} \tag{SN.9}$$

where $sF(t) = f_F(t) \cdot sF(t_{P40})$ and where we assumed that $f_{FB}(t)$ is a tanh function with

$$f_{FB}(t, t_{1/2}, h) = 2^{-h} \cdot \left(1 + \tanh\left[\frac{3}{t_{1/2}} \cdot (t - t_{1/2})\right]\right)^h \tag{SN.10}$$

that models a time-dependent increase in the propensity to form bulbous tips. We had set $h = 1$ and $t_{1/2} = 1000$, such that the rate of bulbous formation peaks at P60–P80, compare Fig. 3B (main manuscript).
Note, for this particular (linear) model, one can also fit $r_3(t)$ and $c_5$ to the marginal distributions of $sB$, $synB$ depicted in Fig. SN.8 and Fig. SN.10, such that $sB \sim \mathcal{P}(\lambda_1)$ and $synB \sim \mathcal{P}(\lambda_2)$ with $\lambda_1 = r_3/(c_4 + c_5)$ and $\lambda_2 = \lambda_1 \cdot c_5/c_4$.

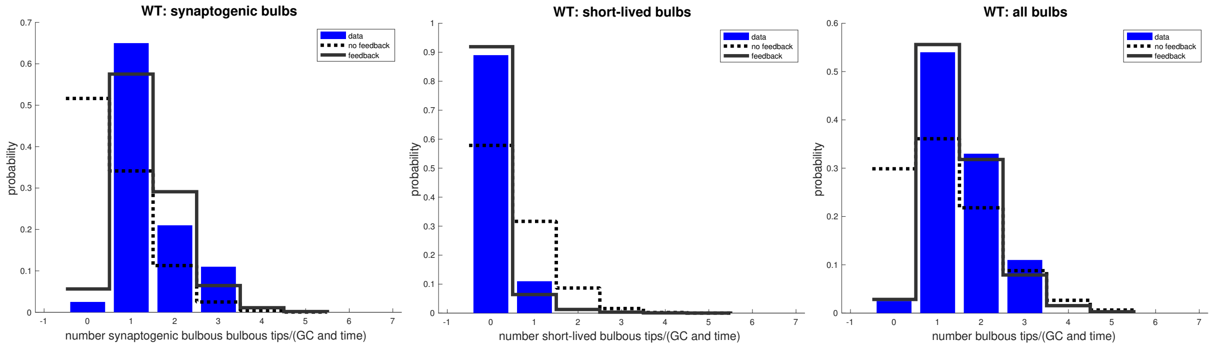


Figure SN.3: **Bulbous tip numbers.** Measured (blue bars) number distribution of bulbous tips at P60 in the wild type. Predicted (dashed black line vs. solid grey line) number distribution of bulbous tips when there is no feedback inhibition vs. in the presence of auto-inhibition.

**Model II: Feedback on bulbous generation.** We followed the analogous procedure as for model I, only that we incorporated a feedback mechanisms into the generator matrix

$$G([i,j],[i-1,j]) = i \cdot c_4, \qquad G([i,j],[i,j-1]) = j \cdot c_6 \tag{SN.11}$$

$$G([i,j],[i+1,j]) = r_3(t) \cdot f_1(i+j, B_{50}), \qquad G([i,j],[i,j+1]) = i \cdot c_5, \tag{SN.12}$$

i.e. $r_3(t)$ is auto-inhibited by the number of bulbous tips present via the function $f_1(B, B_{50})$ [2] (page 247):

$$f_1(B, B_{50}) = \frac{B_{50}}{B_{50} + B},$$

where $B = sB + synB$ denotes the total number of bulbous tips. The resulting fit for the wild type is shown in Fig SN.3, showing that this model can capture the bulbous tip dynamics much better that model I. Essentially, model II results in extremely few non-synaptogenic bulbous tips and guarantees that at least one stabilised (synaptogenic) bulbous tip is present at all times, as observed for the wild type. The biological mechanism behind this feedback could be a general resource limitation and specific allocation of this resource to bulbous tips, such that further bulbous tips cannot be formed.

**Synapse generation, $r_6$.** As mentioned earlier we assumed first-order kinetics and in line with the serial synapse formation model assumed that only one bulbous can generate a synapse at a time, deriving

$$r_6 = c_6 \cdot \min(synB, 1). \tag{SN.13}$$

Parameter $c_6 \approx 1/120$ (min$^{-1}$) was then approximated from the maximum slope of synapse generation presented in Figure 3F (main manuscript).

#### SN.1.3 Wild type model and parameters

The reactions rate/propensities of the stochastic model are given by

$$r_{1,sF}(t) = f_F(t) \cdot c_{1,sF}\,, \qquad r_{2,sF}(sF) = sF \cdot c_{2,sF} \tag{SN.14}$$

$$r_{1,\ell F}(t) = f_F(t) \cdot c_{1,\ell F}\,, \qquad r_{2,\ell F}(F) = \ell F \cdot c_{2,\ell F} \tag{SN.15}$$

$$r_3(t, sF, \ell F, B) = c_3 \cdot (sF + \ell F) \cdot f_1(B, B_{50}) \cdot f_{FB}(t, t_{1/2}, h)\,, \qquad r_4(sB) = c_4 \cdot sB \tag{SN.16}$$

$$r_5(sB) = c_5 \cdot sB\,, \qquad r_6(synB) = c_6 \cdot \min(1, synB) \tag{SN.17}$$

Using the methods explained in the previous sections, we derive parameters depicted in Table SN.2 for the wild type. In summary, we first estimate $c_{2,sF}$, $c_{2,\ell F}$ from the filopodial life time data (Fig. SN.2). Using the mean number of $sF$, $\ell F$ at P40 (Fig. SN.6, upper left panels), we can then estimate $c_{1,sF}$, $c_{1,\ell F}$. Using these parameters and taking the slow-scale dynamics in Fig. 3D (main manuscript) into account, we fit the fifth-order polynomial $f_F(t)$.

From the life times of bulbous tips (Table SN.3), we can estimate $c_4$, which we use together with the number distribution of short-lived and synaptogenic bulbous tips, to estimate $B_{50}$, $c_5$ and $r_3(t)$ in the auto-inhibition model (model II), Fig. SN.3. Using all parameter estimates derived so far and setting $h = 1$, $t_{1/2} = 1000$ (min) in function $f_{FB}(t, t_{1/2}, h)$, we can estimate parameters $c_3$ as outlined in eq. (SN.9).

| | $c_{1,sF}$ | $c_{2,sF}$ | $c_{1,\ell F}$ | $c_{2,\ell F}$ | $c_3$ | $c_4$ | $c_5$ | $c_6$ | $B_{50}$ | $t_{1/2}$ | $h$ |
|---|---|---|---|---|---|---|---|---|---|---|---|
| wt | 3.88 | 0.69 | 1.15 | 0.11 | 0.022 | 1/120$^+$ | 0.1133 | 1/120$^+$ | 0.0282 | 1000 | 1 |
| dlar | 2.63 | 0.69 | 1.49 | 0.11 | 0.0072 | 0.3 | 0.0228 | 1/120$^+$ | $10^{-4}$ | 1000 | 1 |
| liprin-A | 3.12 | 0.69 | 0.99 | 0.11 | 0.0152 | 0.111 | 0.0028 | 1/120$^+$ | 0.363 | 1000 | 1 |
| syd-1 | 2.84 | 0.69 | 1.07 | 0.11 | 0.0321 | 0.169 | 0.0048 | 1/120$^+$ | 1.084 | 1000 | 1 |
| trio | 4.71 | 0.69 | 1.61 | 0.11 | 0.0139 | 0.1311 | 0.1865 | 1/120$^+$ | 0.0231$^*$ | 1000 | 1 |

Table SN.2: Parameter values of the model. All parameters are in units $(\text{min})^{-1}$, except for $t_{1/2}$ (min), $B_{50}$ and $h$ (unit less). The fifth-order polynome $f_F(t) = \max(0, \sum_{i=0}^{5} p_i \cdot t^i)$ has coefficients $p_5 = -2.97 \cdot 10^{-14}, p_4 = 3.31 \cdot 10^{-13}, p_3 = -1.29 \cdot 10^{-9}, p_2 = 2.06 \cdot 10^{-6}, p_1 = -1.45 \cdot 10^{-3}$ and $p_0 = 1$ $(\text{min}^{-1})$. $^+$ could not be determined from data and was set to 1/120 minutes (almost all wild type bulbous tips eventually become synaptogenic).$^*$Note that the trio feedback mechanisms is modelled slightly different as outlined below.

| WT | DLar | LiprinA | Syd1 | Trio |
|---|---|---|---|---|
| 120(-)$^+$ | 3.3(3) | 9(9.6) | 5.9(6.4) | 7.6(6.8) |

Table SN.3: Average (standard deviation) life time of short-lived unstable bulbous tips $sB$ (min) at P60. $^+$ could not be determined from data and was set to 120 minutes.

##### SN.1.4 Simulation of growth cone retraction.

In Fig. 6 (main manuscript), we depict the simulated probability of growth cone retraction up to time $T$, $P_{\text{retract}}(T)$. The probability of growth cone retraction was computed as

$$P_{\text{retract}}(T) = 1 - \prod_{i=0}^{N} P_{\text{no-retract}}(i) \tag{SN.18}$$

where $P_{\text{no-retract}}(i)$ denotes the probability not to retract in the $i$-th time interval which is computed by

$$P_{\text{no-retract}}(i) = e^{-\Delta t \cdot r_0 \cdot f_{\text{retract}}(F(i),B(i),S(i),w,n_{\text{stab}})} \tag{SN.19}$$

where $r_0$ is the basal rate of retraction, $\Delta t$ is the duration of the $i$-th time interval and $F(i), B(i), S(i)$ are the number of filopodia, bulbous tips and synapses during that time interval. $n_{\text{stab}}$ is the 'minimal stabilization number' and $w$ are the user defined weights, such that:

$$f_{\text{retract}}(F(i), B(i), S(i), w, n_{\text{stab}}) = 0.5 \cdot \left(1 + \tanh\left[\frac{3}{n_{\text{stab}}} \cdot (n(i) - n_{\text{stab}})\right]\right) \tag{SN.20}$$

with $n(i) = w_F \cdot (sF + \ell F) + w_B \cdot (sB + synB) + w_S \cdot S$ being the weighted sum of filopodia, bulbous tips and synapses affecting (preventing) retraction.

##### SN.1.5 Effect of knock-out mutants.

The *life time* of short and long-lived filopodia were not markedly different as shown in Table SN.4 and hence rates $c_{2,sF}$, $c_{2,\ell F}$ were set equal in all mutants and the wild type. However, the *number* of short- and long-lived filopodia were different between wild type in all knock-out mutants, Table SN.5.We modelled these differences by estimating mutant-specific rates $c_{1,sF}$, $c_{1,\ell F}$ as shown in Table SN.2 above.
A striking observation is that we observe two populations of bulbous tips in the mutants (see Fig. SN.4): One population has a short life time (transient), whereas the other one seems to be stable as shown in Table SN.3. Consequently, parameters $c_4$ were set to the inverse of the mutant-specific bulbous tip life times depicted in Table SN.3.

***Trio*-Model: Feedback on bulbous stabilisation:** The *trio* mutant shows somewhat different dynamics to all other mutants and with respect to the wild type: Unlike the wild type, *trio* has both a population of short-lived as well as a population of long-lived bulbous filopodia. Moreover, unlike the other mutants, at least one stabilized (synaptogenic) bulbous tip is present at all times. Consequently, the presence of bulbous tips does not seem to down-regulate the *initiation* of new (short-lived) bulbous tips (as in the wild type), compare Figures. SN.9 and SN.7. Moreover, the distribution of *stabilized* bulbous tips cannot be explained by simple/linear dynamics as in the case of syd-1. Rather, it seems that the presence of stabilized (synaptogenic) bulbous tips may down-regulate bulbous tip *stabilization*. The mechanistic reason for this could be a general resource limitations for factors *stabilizing* bulbous tips. This resource-limitation may not be observable in other mutants, which have very few stabilized bulbous tips at any time instance (syd-1, lar, liprin-alpha). Moreover, the observed resource-limitation in *trio* may also not be observed in the same way in the wild type, because unlike in trio, essentially all bulbous

tips in the wild type are stable.
This observation prompted us to assume a strong auto-inhibitory feedback mechanisms of synaptogenic bulbous tips on their own production.
This generator for this model is as follows

$$G([i, j], [i-1, j]) = \quad i \cdot c_4, \qquad G([i, j], [i, j-1]) = j \cdot c_6 \tag{SN.21}$$

$$G([i, j], [i+1, j]) = \quad r_3(t), \qquad G([i, j], [i, j+1]) = i \cdot c_5 \cdot f_1(j, B_{50}), \tag{SN.22}$$

with parameters stated in Table SN.2.

#### SN.2 Raw data

##### SN.2.1 Bulbous life time distribution

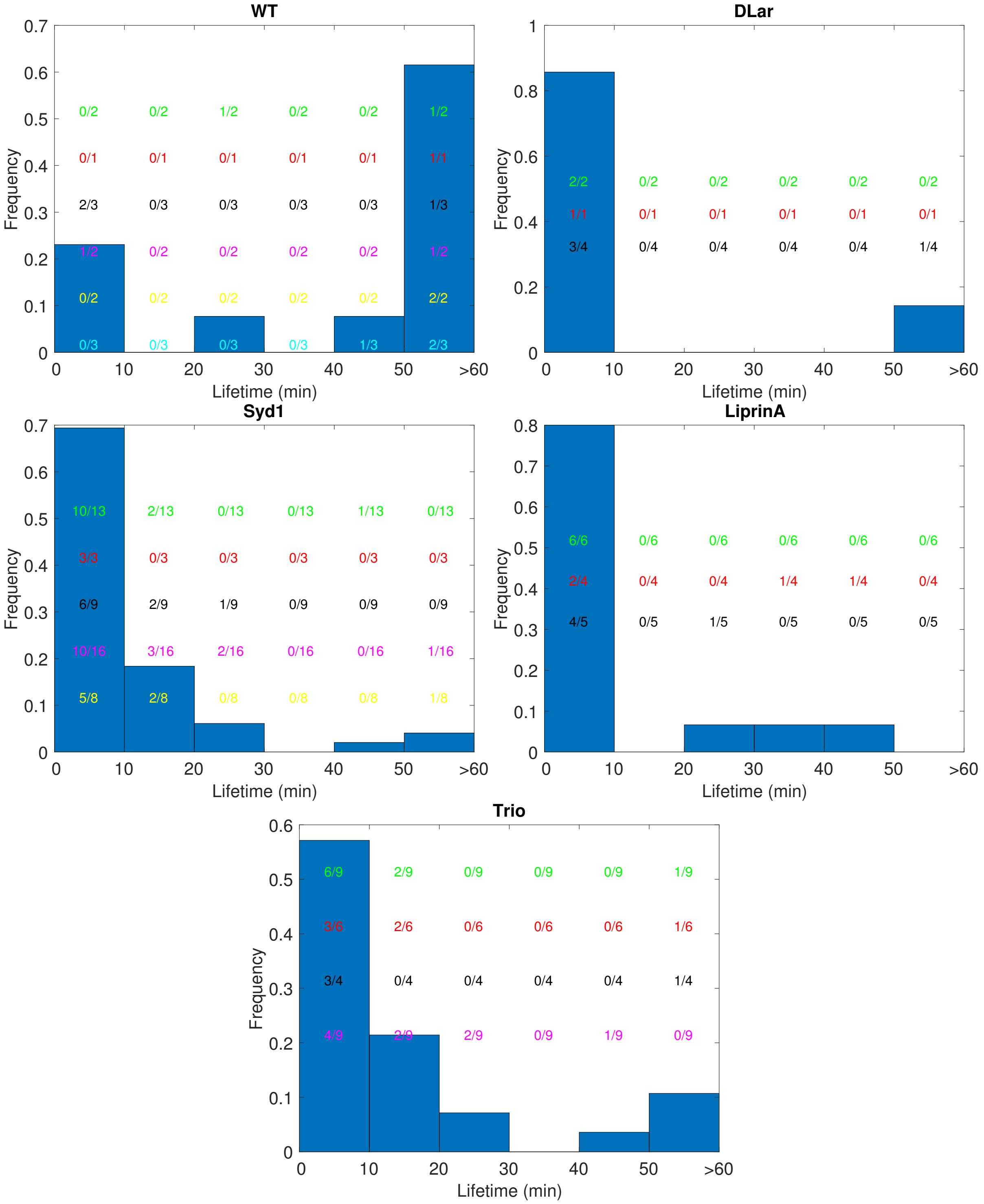


Figure SN.4: **Bulbous life time distribution.** All mutants show a population of short-lived bulbous tips (left bars) that is almost absent in the wild type. Numbers show the counts in each life time category for different growth cones.

#### SN.2.2 Filopodia numbers

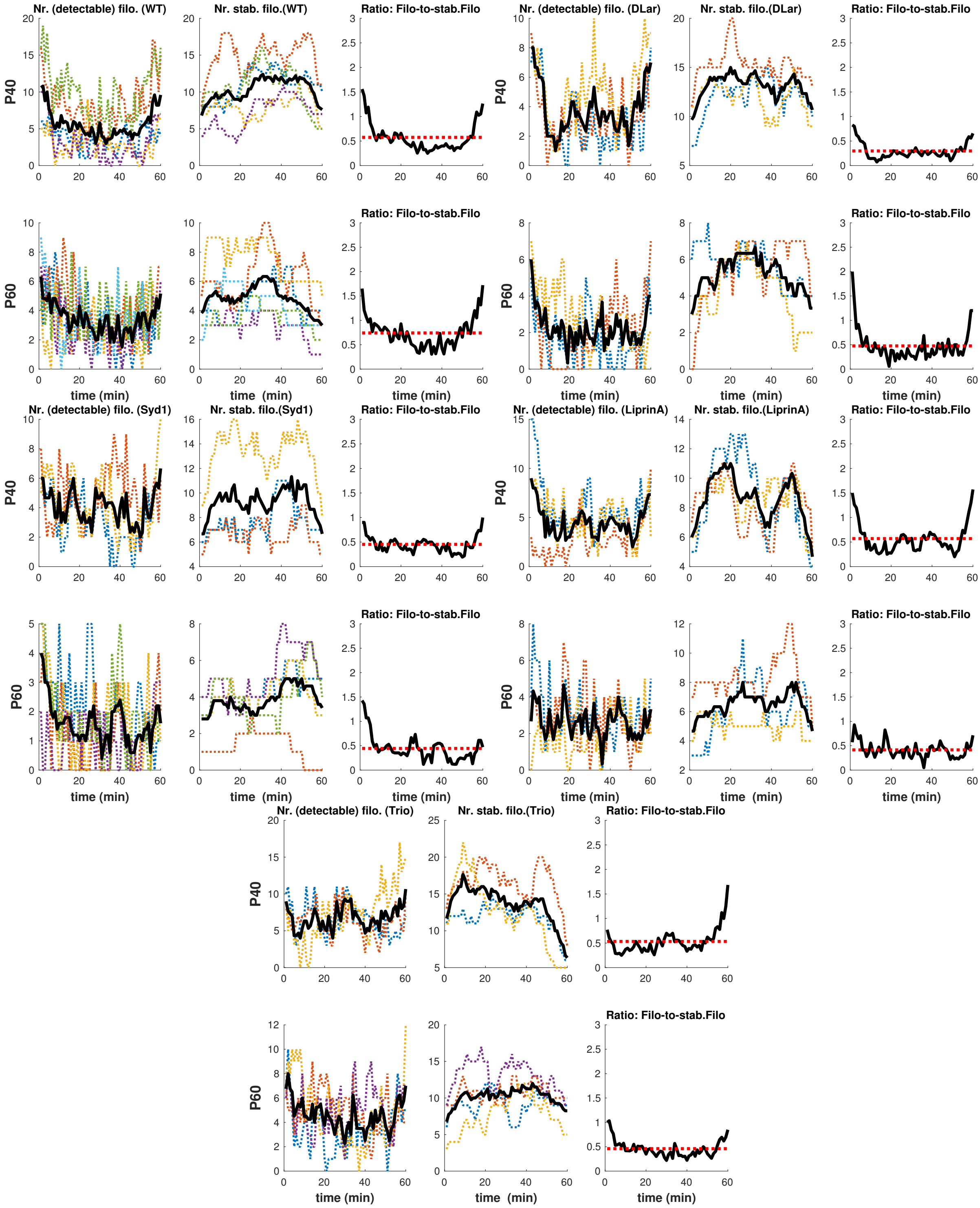


Figure SN.5: **Numbers of filopodia per time instance.** Number distribution of short-lived and stabilized filopodia and their ratio at P40 and P60 respectively in the different mutants. Dashed lines indicate different growth cones, while the solid black lines show the average (over all growth cones) number of filopodia per time instance.

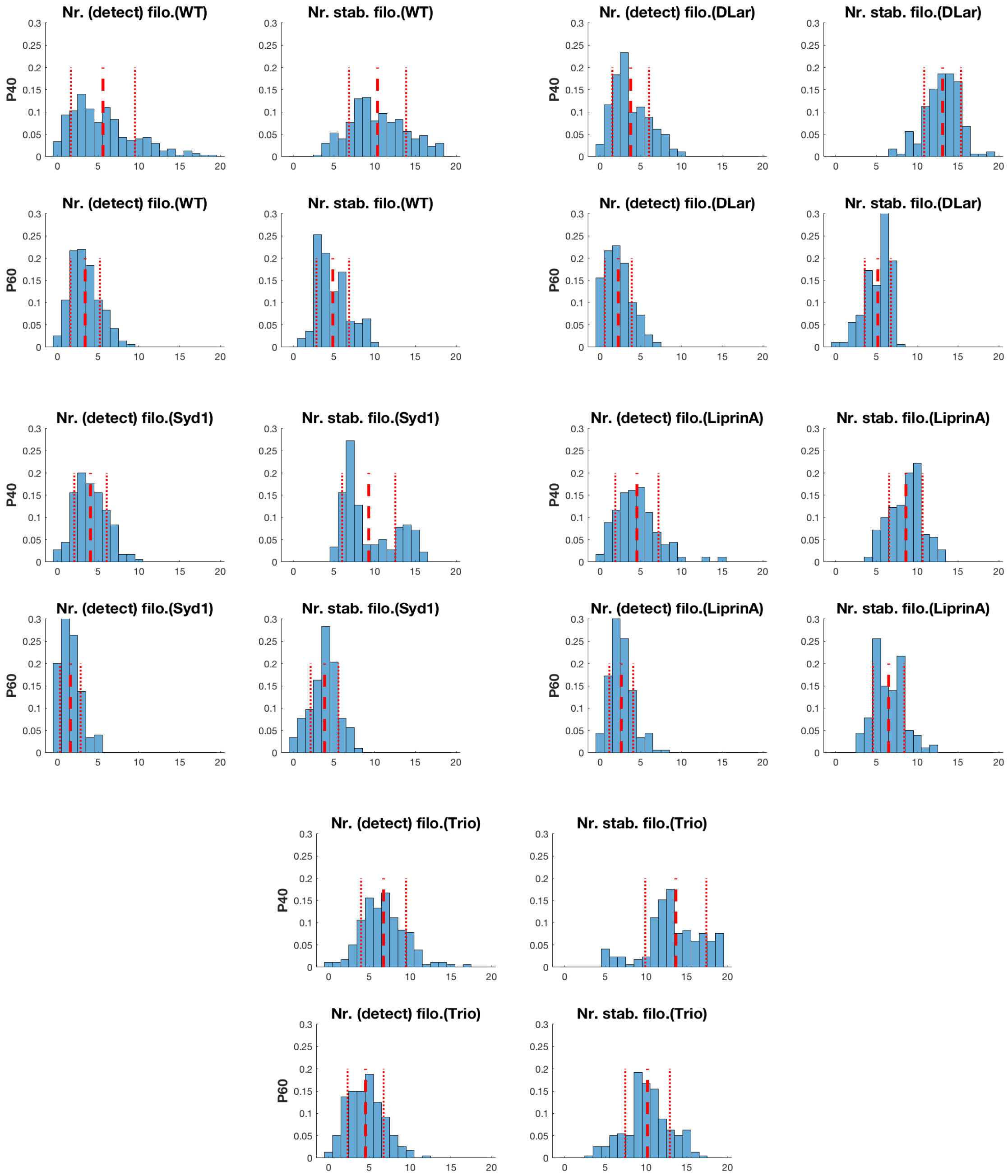


Figure SN.6: **Number distribution of short-lived and stabilized filopodia at P40 and P60 respectively in the different mutants.** We averaged over all growth cones and over time (over the 60minutes fast recordings at P40 and P60 respectively). Thick and thin dashed vertical lines mark the mean number ± one standard deviation.

#### SN.2.3 Transient bulbous numbers

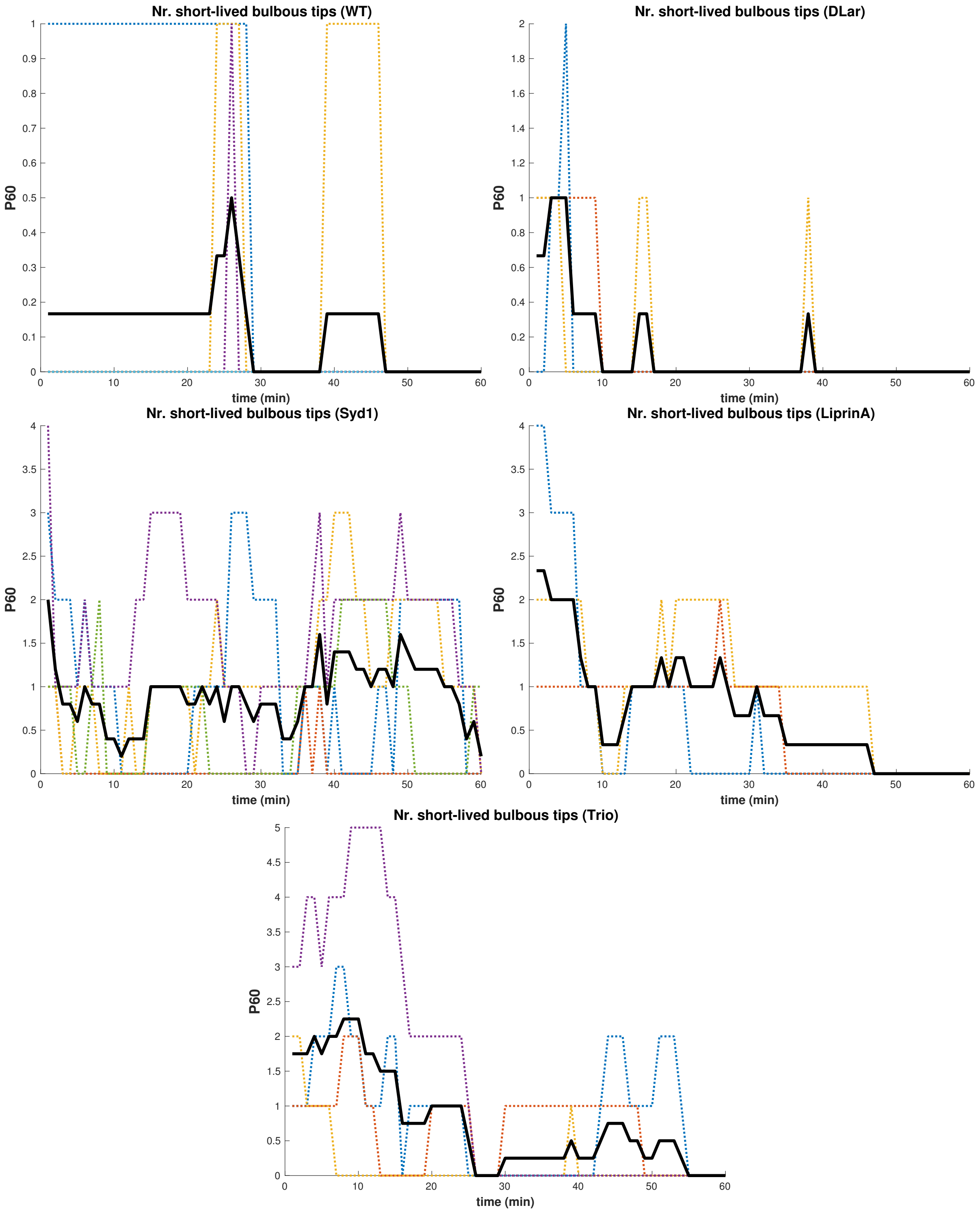


Figure SN.7: **Number of transient bulbous tips per time instance** at P60 per growth cone in the different mutants. Solid black line marks the average number per time instance (averaged over all growth cones) and dashed lines show the number for individual growth cones.

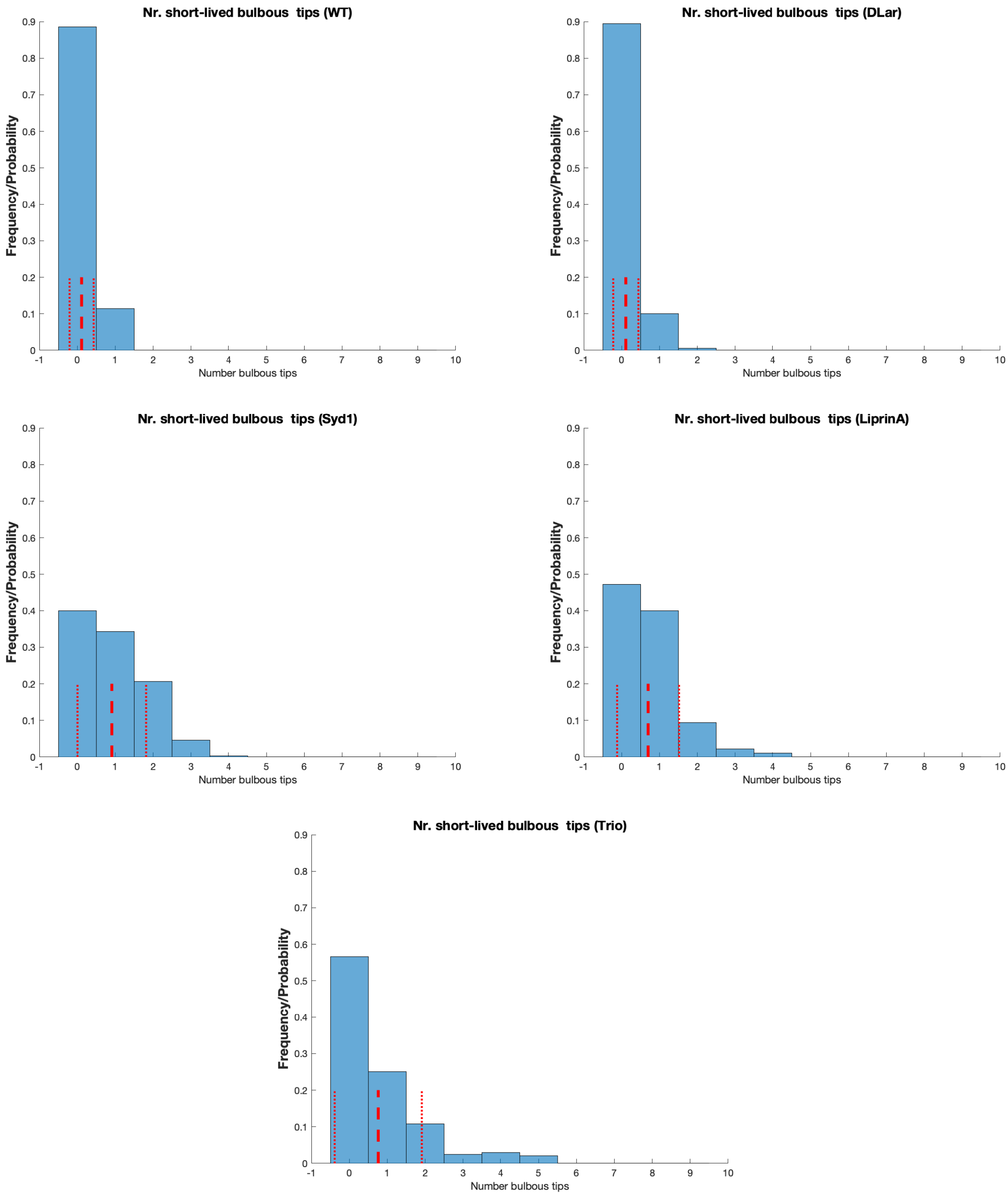


Figure SN.8: **Number of transient bulbous tips** at P60 averaged over all growth cones and over time in the different mutants. Dashed red line marks the average number per time instance ± one standard deviation.

##### SN.2.4 Stabilized bulbous numbers

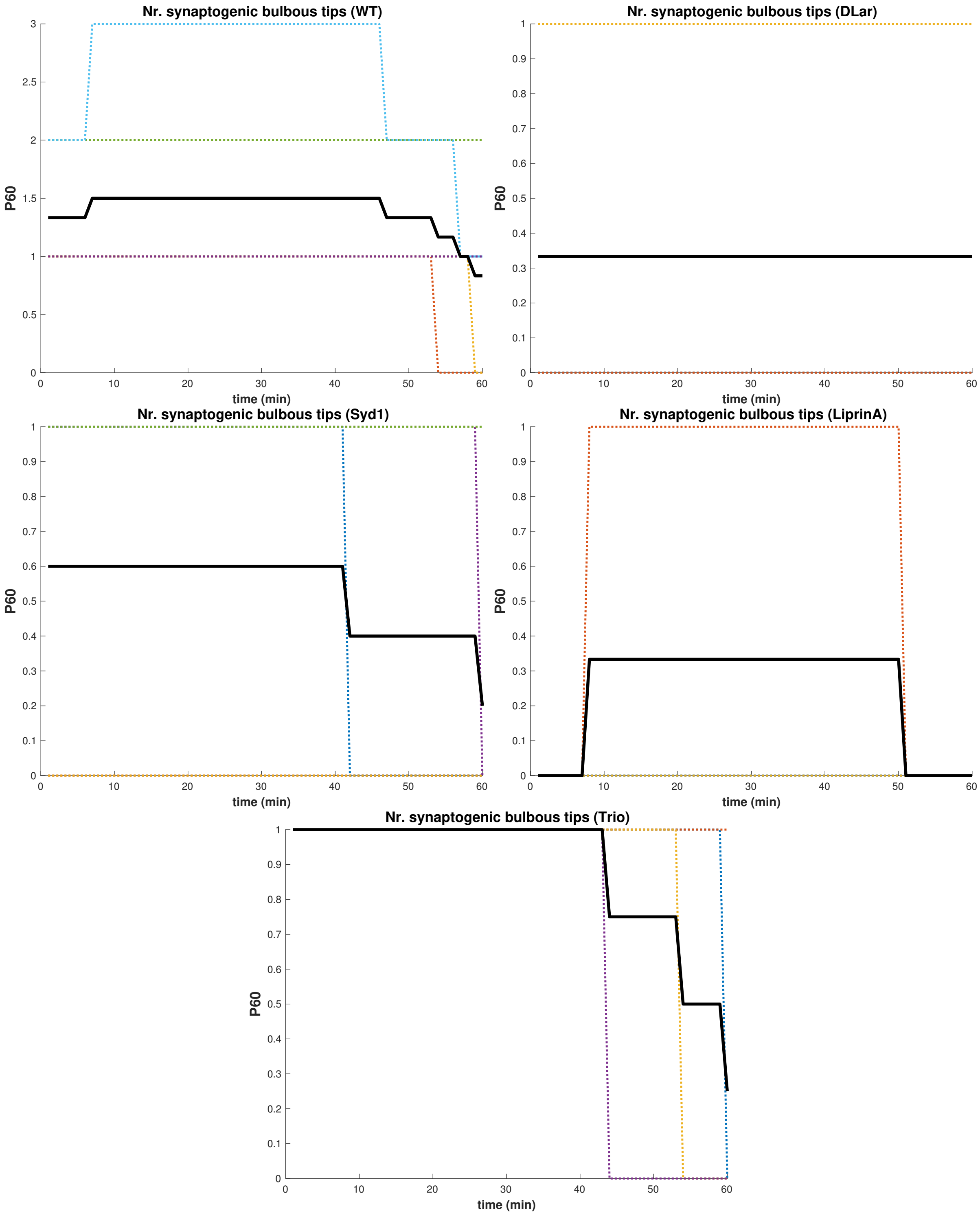


Figure SN.9: **Number of stabilized bulbous tips per time instance** at P60 per growth cone in the different mutants. Solid black line marks the average number per time instance (averaged over all growth cones) and dashed lines show the number for individual growth cones

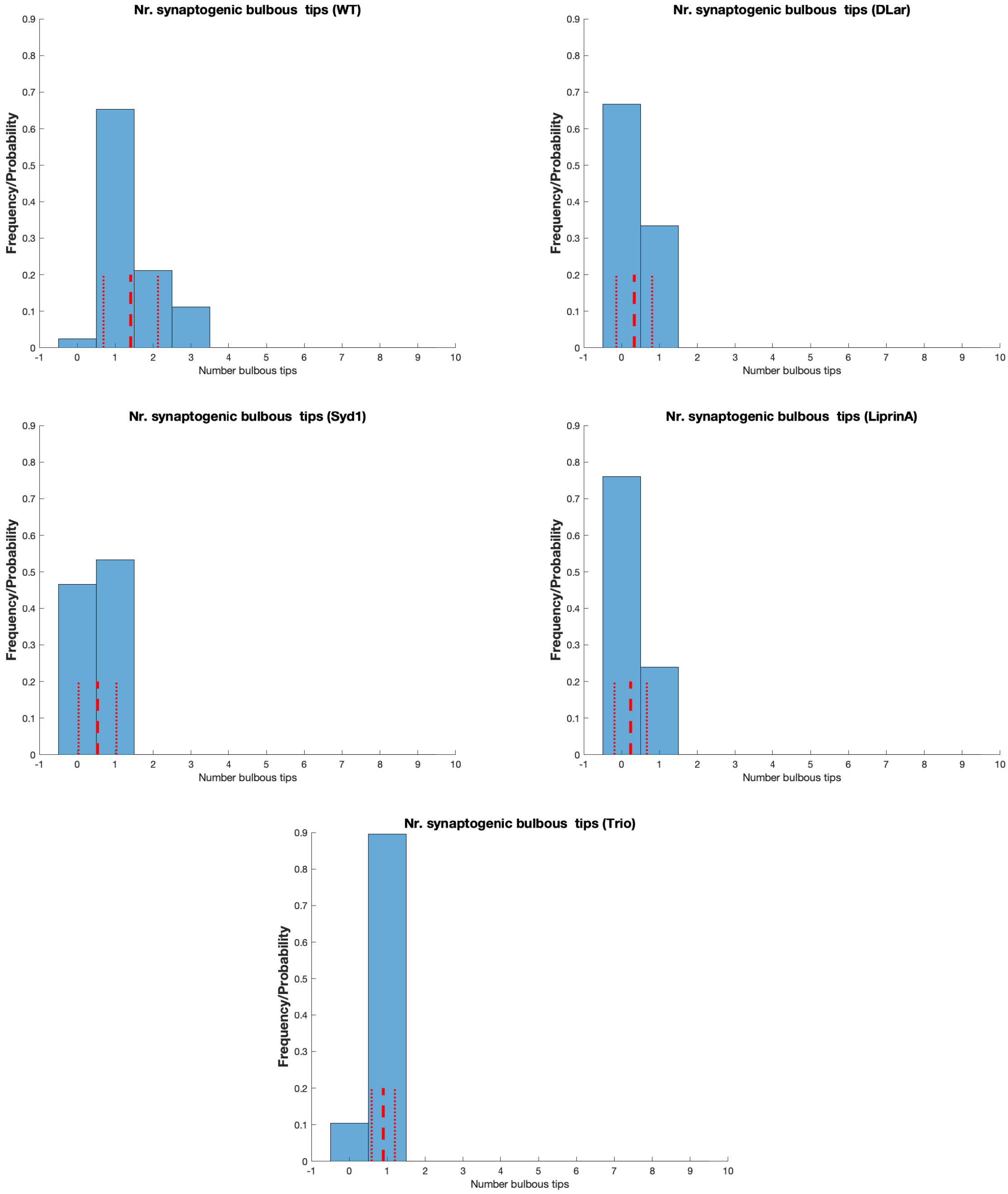


Figure SN.10: **Number of stabilized bulbous tips** at P60 averaged over all growth cones and over time in the different mutants. Dashed red line marks the average number per time instance ± one standard deviation.

#### SN.2.5 All bulbous numbers

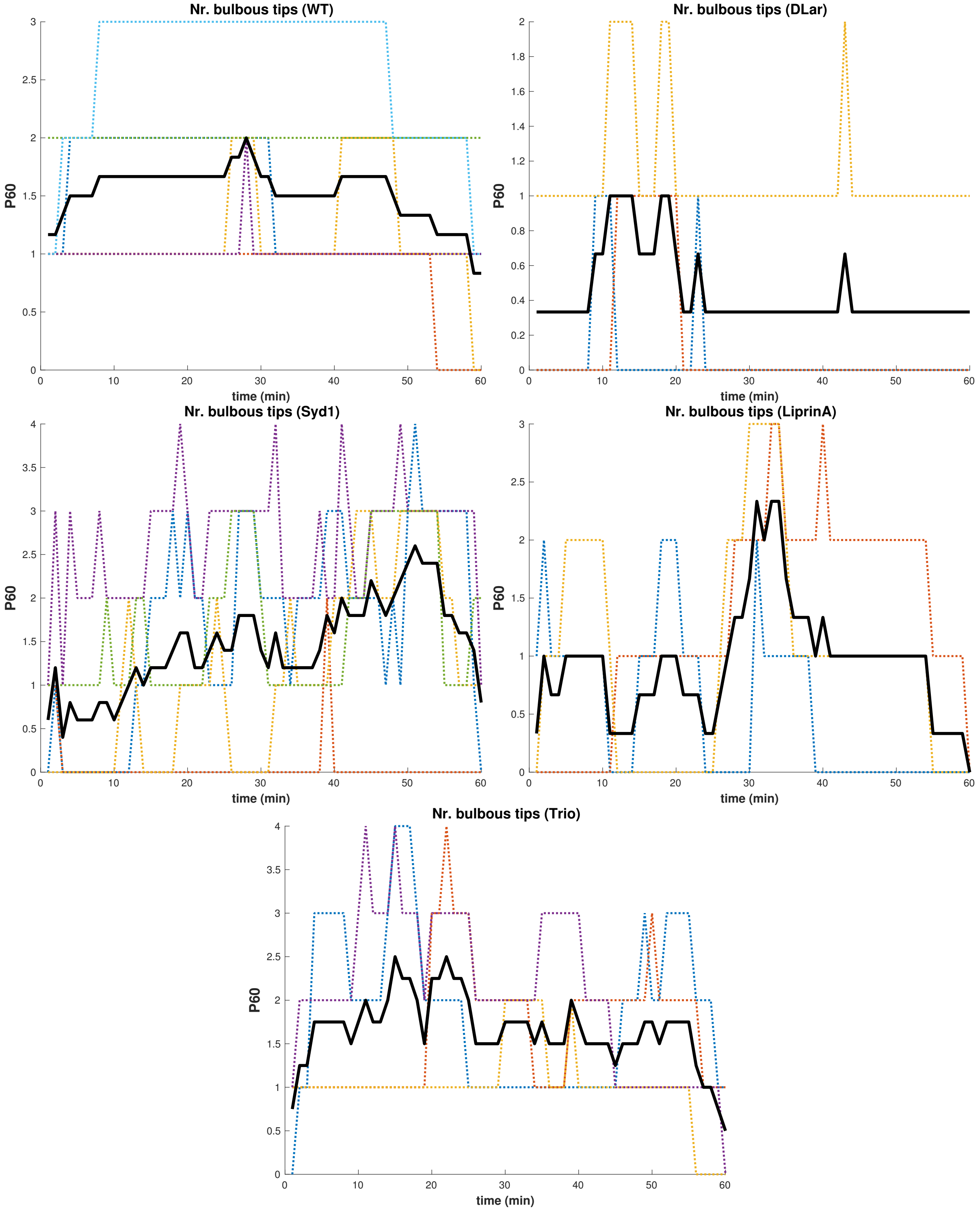


Figure SN.11: **Total number of bulbous tips (stabilized + transient) per time instance** at P60 per growth cone in the different mutants. Solid black line marks the average number per time instance (averaged over all growth cones) and dashed lines show the number for individual growth cones.

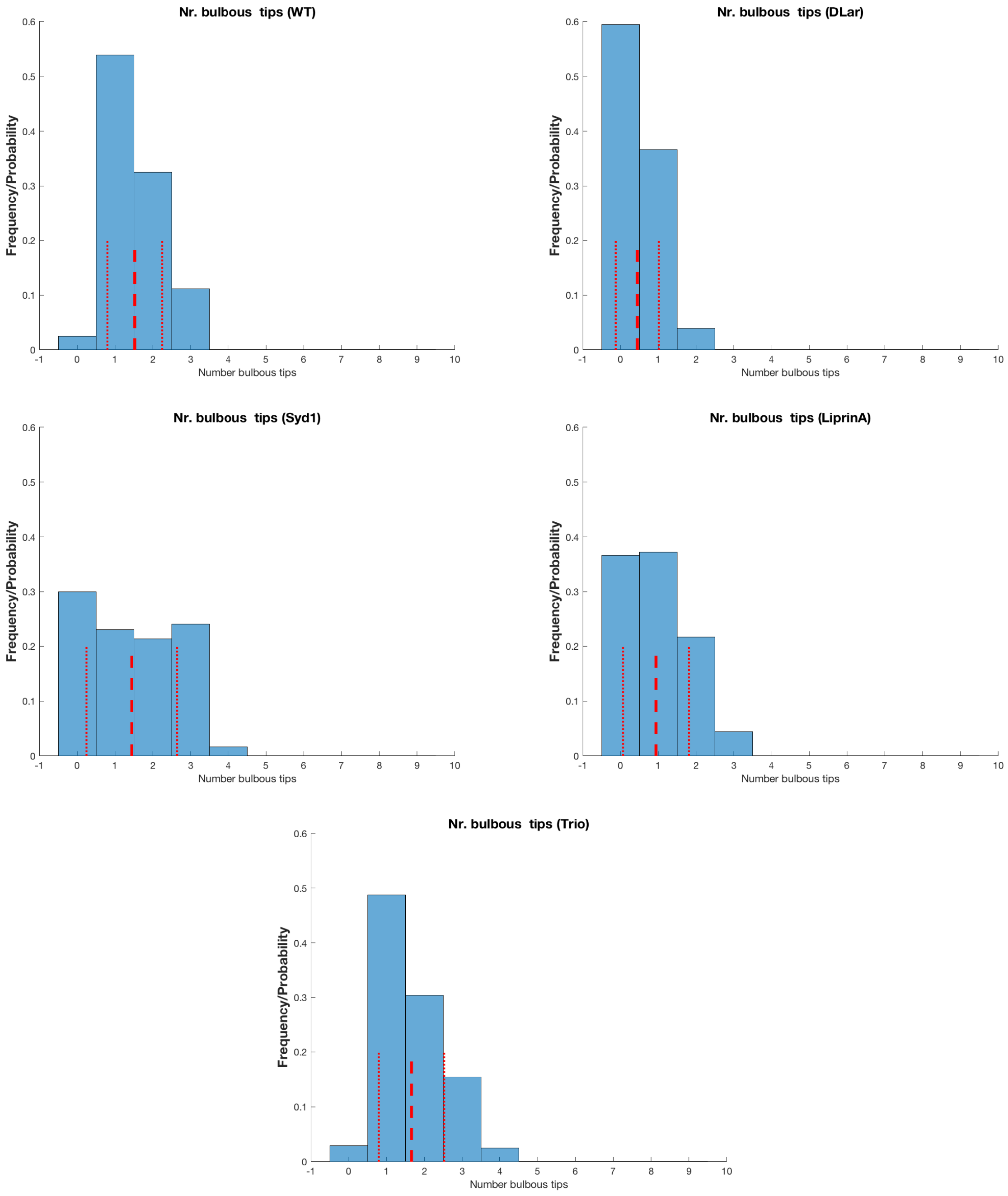


Figure SN.12: **Total number of bulbous tips** (transient + stabilized) at P60 averaged over all growth cones and over time in the different mutants. Dashed red line marks the average number per time instance ± one standard deviation.

##### SN.2.6 Filopodia life times

| | short-lived | | | long-lived | | |
|---|---|---|---|---|---|---|
| Mutant | P40 | P60 | P40 and P60 | P40 | P60 | P40 and P60 |
| WT | 2.4(1.7) | 1.9(1.4) | 2.2(1.6) | 18(13) | 23(18) | 20(15) |
| DLar | 2.7(1.7) | 2.3(1.5) | 2.5(1.6) | 23(17) | 19(15) | 22(16) |
| LiprinA | 2.6(1.9) | 2.3(1.6) | 2.5(1.8) | 18(13) | 20(15) | 19(14) |
| Syd1 | 2.3(1.6) | 2.2(1.7) | 2.3(1.7) | 18(13) | 23(16) | 20(14) |
| Trio | 2.3(1.7) | 2.6(1.8) | 2.5(1.8) | 19(12) | 20(15) | 20(14) |

Table SN.4: Average (standard deviation) life time of filopodia (min) that were classified as short-lived vs. long-lived based on the 8min criterium.

##### SN.2.7 Filopodia numbers

| | short-lived | | long-lived | |
|---|---|---|---|---|
| Mutant | P40 | P60 | P40 | P60 |
| WT | 5.6(3.9) | 3.4(1.8) | 10(3.5) | 4.9(2) |
| DLar | 3.8(2.3) | 2.2(1.7) | 13(2.3) | 5.2(1.6) |
| LiprinA | 4.5(2.7) | 2.6(1.5) | 8.6(2.1) | 6.5(1.9) |
| Syd1 | 4.1(2) | 1.6(1.3) | 9.3(3.3) | 3.8(1.7) |
| Trio | 6.8(2.8) | 4.5(2.2) | 14(3.7) | 10(2.7) |

Table SN.5: Average (standard deviation) numbers of short-lived and long-lived filopodia per time instance

[2] E Klipp, R Herwig, A Kowald, C Wierling, and H Lehrach. *Systems Biology in Practice*. Wiley-VCH, 2006.

**Supplementary Figure Legends Özel et al.**

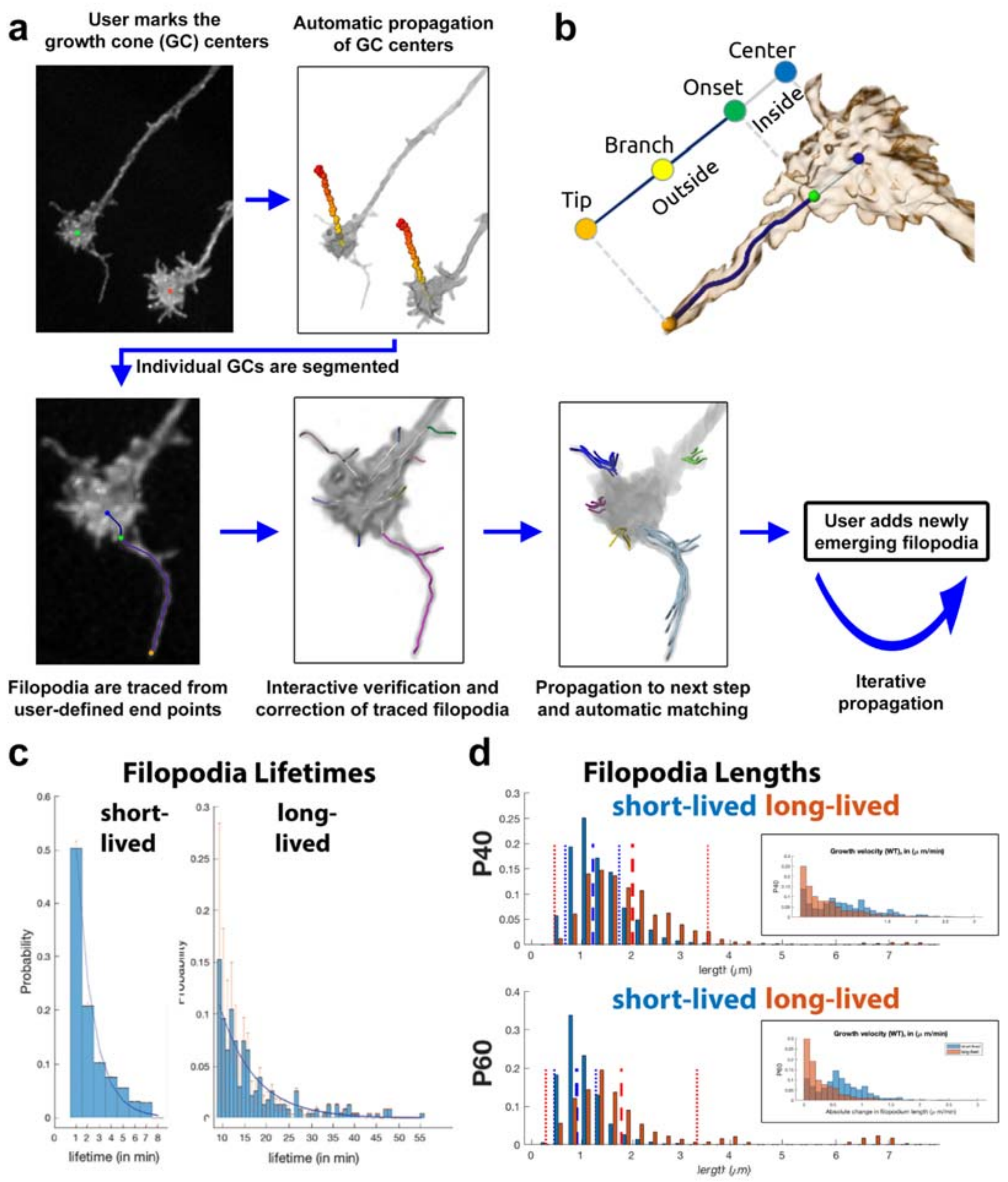


**Supplementary Figure 1: Filopodia quantification. (a)** The user selects the growth cone (GC) for further processing by marking the centers (color-coded) in the first timestep. Their successors are automatically detected in the following time steps. To process the GCs one at a time, the user adds a new filopodium by interactively specifying the tip (orange). The path to the GC center (blue) is then automatically traced. The filopodia onset (green) is set to the location on the path where the intensity profile in a plane orthogonal to the path changes from Gaussian to non-Gaussian. The onset point divides the path into an inside (blue) and an outside (purple) part, the latter being the actual filopodium. A different track ID (color-coded) is assigned to each filopodium. **(b)** A neuronal growth cone is represented as a skeleton graph (tree). One branch of the tree extended from the GC center (blue) to the filopodia tip (orange), passing through the onset location (green) and potentially branching nodes (yellow). The part of the path between tip and onset ("Outside") is the actual filopodium. **(c)** All filopodial lifetimes can be described with two exponential distributions, one for short-lived and one for long-lived filopodia. **(d)** Filopodial length distributions can be described with separate distributions for short-lived and long-lived filopodia, both for P40 and P60.

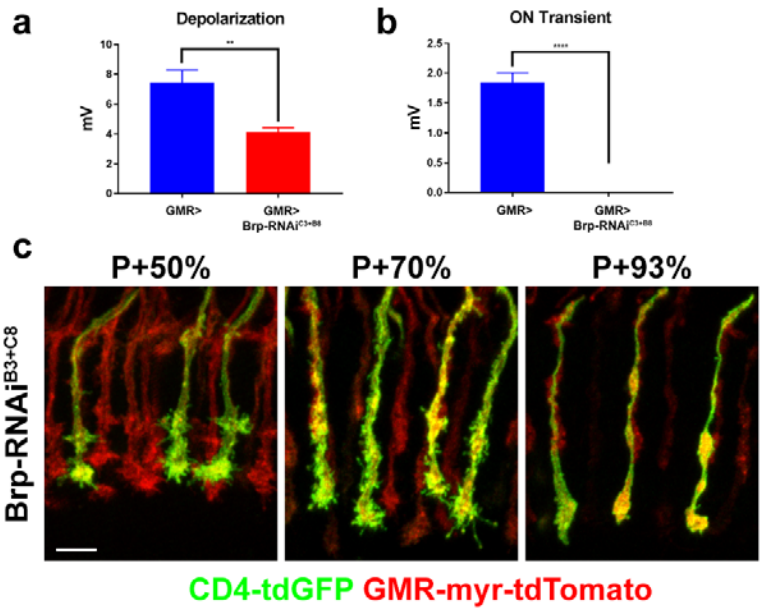


**Supplementary Figure 2: Bruchpilot is not required for normal development of R7 axons.** ERG recordings showing the level of **(a)** depolarization ($p = 0.0014$) and **(b)** ON Transient ($p<0.0001$) from eyes that express only GMR-Gal4 and those that co-express two RNAi constructs against the *brp* gene, Brp-RNAi$^{B3}$ and Brp-RNAi$^{C8}$. Error bars denote SEM. **(c)** Sparsely generated R7 clones during pupal development are labeled with CD4-tdGFP and co-express the two RNAi constructs. All photoreceptors are marked with myr-tdTomato. Scale bar: 5 µm.

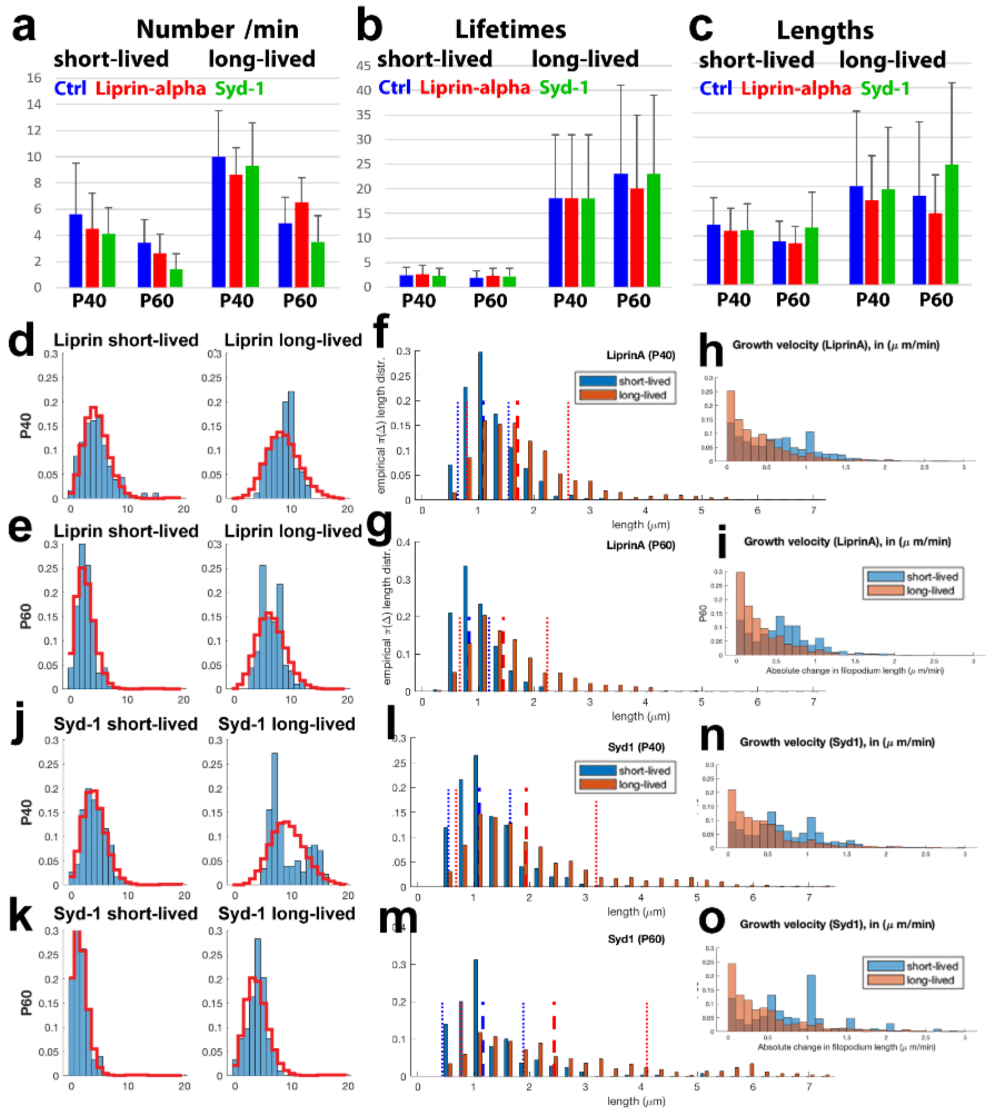


**Supplementary Figure 3: Number, lifetime, length and velocity statistics for filopodia in liprin-alpha and syd-1 mutants.** **(a-c)** Numbers, lifetimes (min) and lengths of short-lived and long-lived filopodia at both P40 and P60 are not statistically different in ctrl, *liprin-α* and *syd-1*. **(d-e)** Numbers of short-lived (left) and long-lived (right) filopodia in the *liprin-α* mutant resemble Poisson distributions. **(f-i)** Lengths and velocity distributions of short-lived and long-lived filopodia in the *liprin-α* mutant resemble Poisson distributions.

**(j-k)** Numbers of short-lived (left) and long-lived (right) filopodia in the *syd-1* mutant resemble Poisson distributions, except for some long-lived filopodia at P40. **(l-o)** Lengths and velocity distributions of short-lived and long-lived filopodia in the *syd-1* mutant resemble Poisson distributions except for a few particularly long, long-lived filopodia.

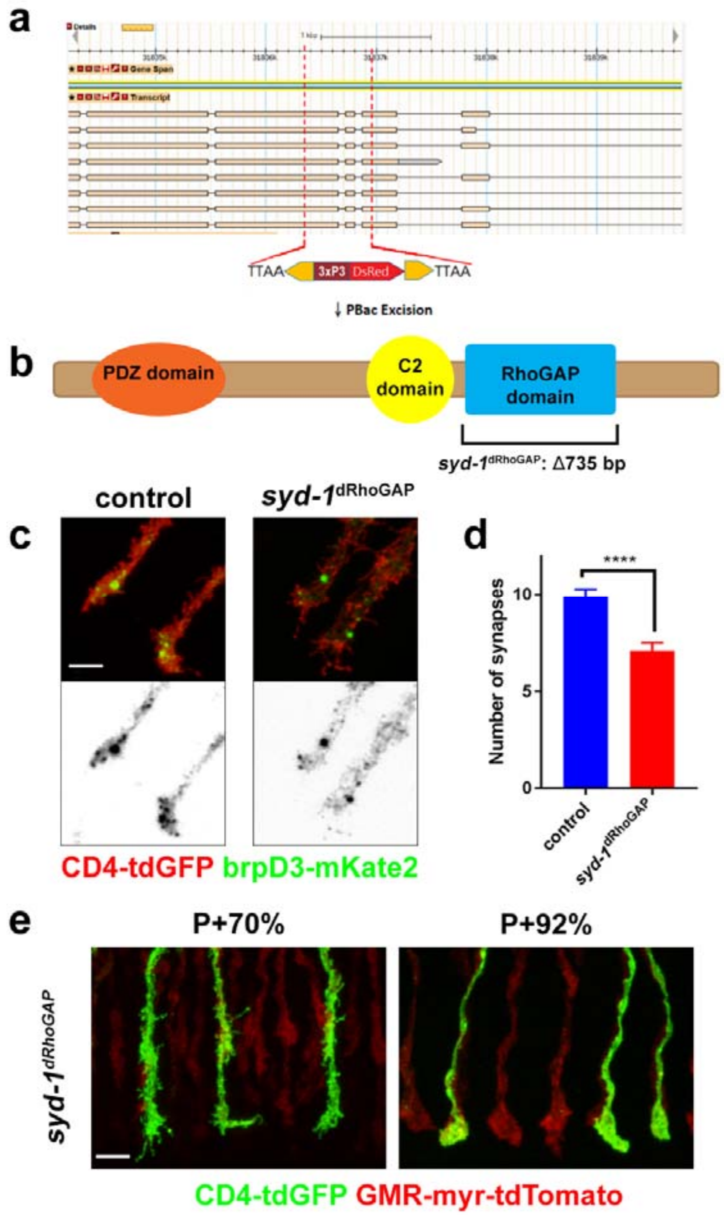


**Supplementary Figure 4: RhoGAP domain of Syd-1 is not required for axonal development in R7 axons. (a)** CRISPR-mediated knock-in of the Scarless construct into the *syd-1* locus. Upon PBac excision of the DsRed casette **(b)** RhoGAP domain of Syd-1 is deleted completely and specifically, leaving the rest of the protein intact. **(c)** Presynaptic punctae at P+70% in sparsely generated *syd-1*$^{dRhoGAP}$ R7 terminals and FRT82B controls. **(d)** Quantification of b (n= 45 and 32, $p<0.0001$) **(e)** Sparsely generated *syd-1*$^{dRhoGAP}$ clones marked with CD4-tdGFP and all photoreceptors marked with myr-tdTomato. Mutant axons appear normal at P+70% and 92%. Scale bar: 5 µm.

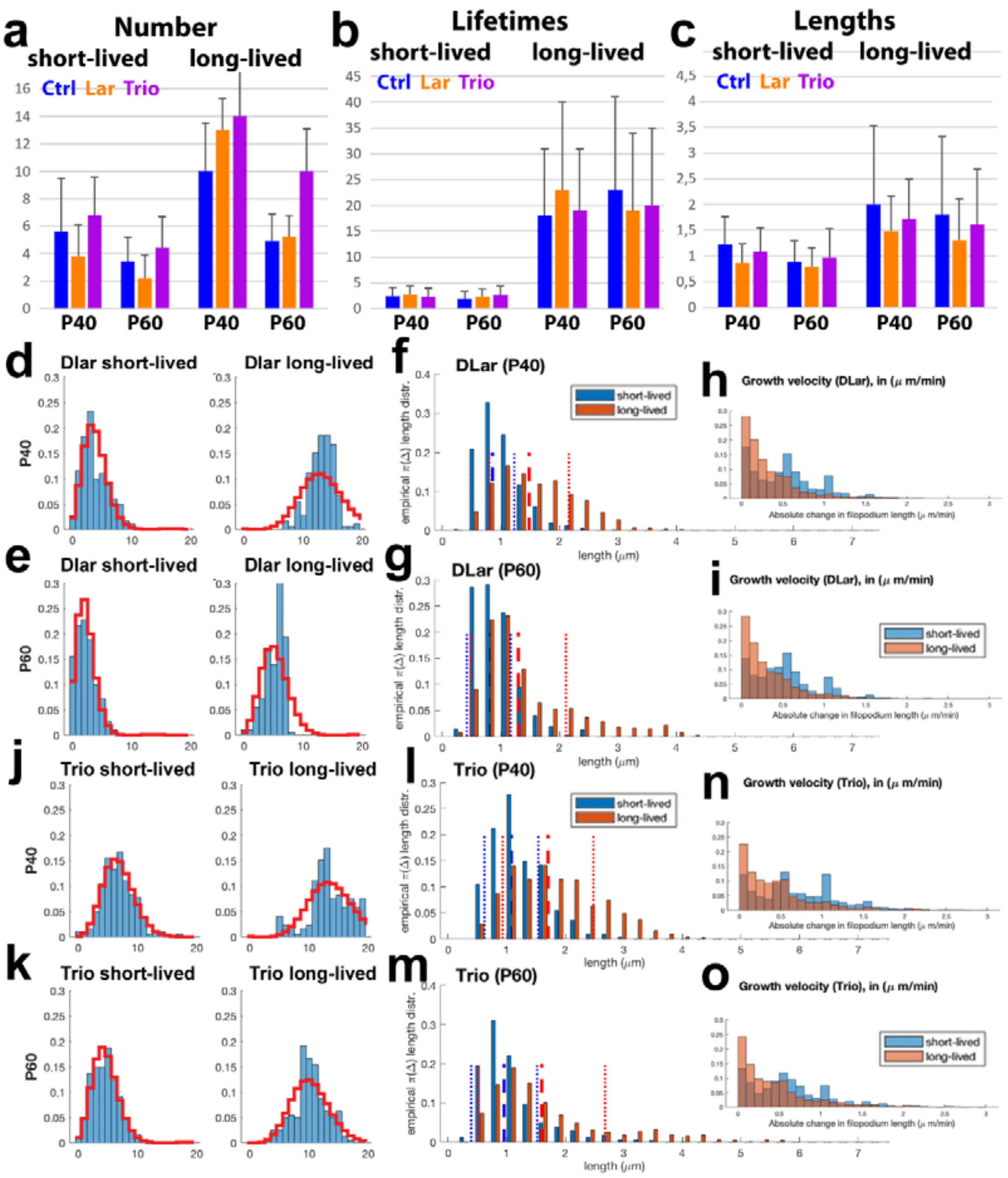


**Supplementary Figure 5: Number, lifetime, length and velocity statistics for filopodia in lar and trio mutants. (a-c)** Numbers, lifetimes and lengths of short-lived and long-lived filopodia at both P40 and P60 are not statistically different in ctrl, *lar* and *trio*, except for mild increases of filopodia numbers in trio at P60. **(d-e)** Numbers of short-lived (left) and long-lived (right) filopodia in the *lar* mutant resemble Poisson distributions. **(f-i)** Lengths and velocity distributions of short-lived and long-lived filopodia in the *lar* mutant resemble separate Poisson distributions. **(j-k)** Numbers of short-lived (left) and long-lived (right) filopodia in the *trio* mutant resemble Poisson distributions. **(l-o)** Lengths and velocity distributions of short-lived and long-lived filopodia in the *trio* mutant resemble separate Poisson distributions.

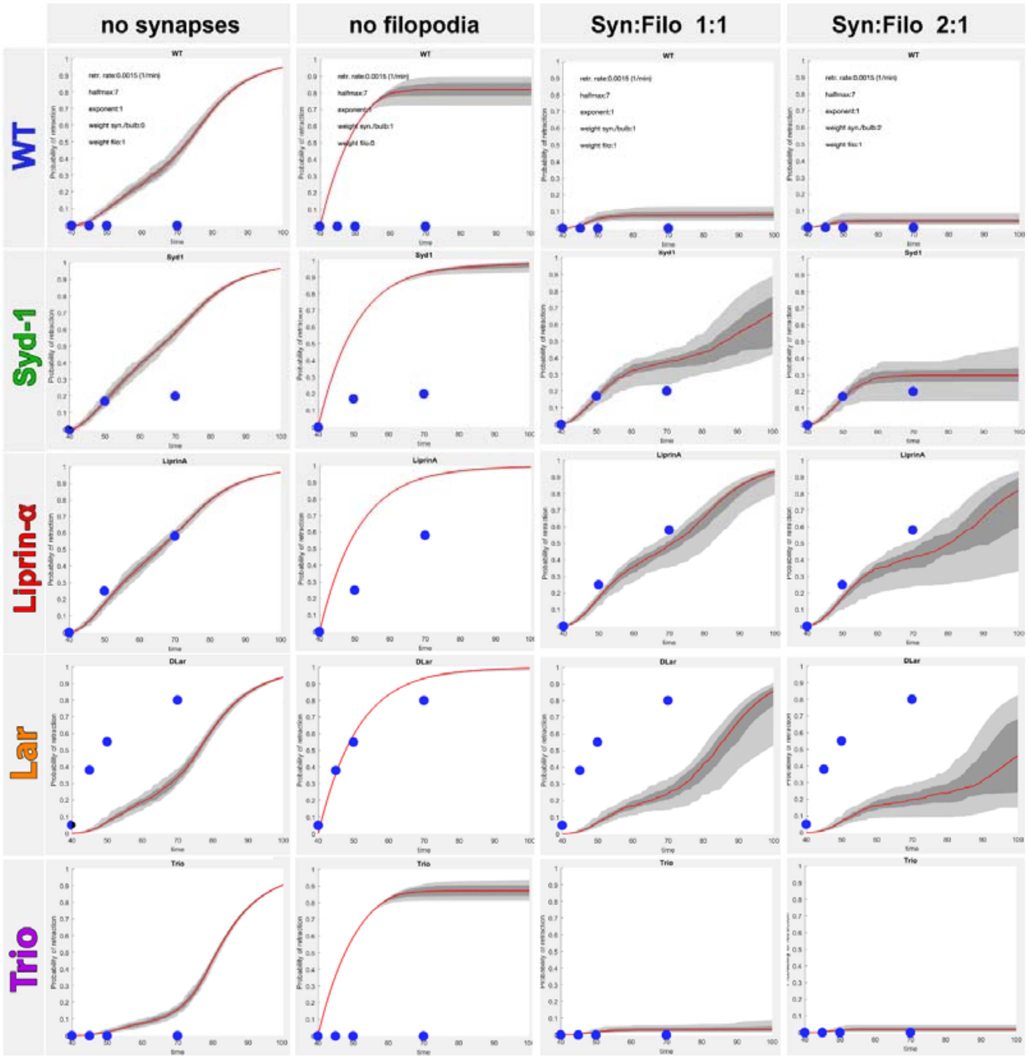


**Supplementary Figure 6: Modeling of retraction probabilities as a function of stabilization through filopodia and synapses**

First column (left): Simulation of retractions if synapses do not contribute to axon terminal stabilization (but only filopodia stabilize terminals). All mutants would exhibit retractions after P50 if synapses do not contribute to stabilization.

Second column: Simulation of retractions if filopodia do not contribute to axon terminal stabilization (but only synapses stabilize terminals). Retractions would be similar in all mutants due to absence of synapses before P50.

Third column: Simulation of equally weighted filopodia and synapses contributing to axon terminal stabilization. Wild type and *trio* exhibit no retractions, as declining filopodia numbers are compensated for by increasing synapse numbers. lar, *liprin-α* and *syd-1* exhibit retractions with different kinetics based on defects in synapse formation.

Fourth column (right): Simulation of stabilization weighted 2:1 to synapses over filopodia only mildly changes retraction dynamics and matches closely the observed retraction kinetics in all mutants except *lar*. *lar* is best matched if filopodia do not contribute to stabilization (second column), suggesting a loss of filopodial adhesion as well.

blue discs: measured retraction values.

**Table 1: Measured rates of bulb dynamics at P60**

| | r3 | r2B | E[f1] | r4 | r5 | Avg. bulbs |
|---|---|---|---|---|---|---|
| **WT** | 0,0222 (0,0172) | 0,1636 | 0,0768 | 0,0083 | 0,0117 | 1,5300 |
| **Liprin** | 0,0833 (0,0167) | 0,0949 | 0,8347 | 0,0778 (0,0255) | 0,0019 | 0,9526 |
| **Syd1** | 0,16 (0,0847) | 0,2049 | 0,7737 | 0,1533 (0,0767) | 0,0044 | 1,4296 |
| **Dlar** | 0,0333(0,0167) | 0,0576 | 0,6736 | 0,0333 (0,0167) | 0,0027 | 0,4477 |
| **Trio** | 0,1125 (0,0438) | 0,1381 | 1,0000 | 0,1 (0,0408) | 0,0141 | 1,6331 |

r3: measured rate of bulb formation, contains r2B * f1, unit: 1/min
r2B: propensity to form bulbs, cannot be measured, because feedback f1 reduces r2B, shown is the only possible fit of r2B, unit: 1/min
f1: negative feedback on bulb formation, cannot be measure, see r5, shown is the only possible fit of the data (r2B
r4: measured rate of bulbs disappearance, unit: 1/min
r5: measured rate of bulb stabilization, unit: 1/min
Avg. bulbs: average number of bulbs per time instance (min) over an hour (P60)
In blue: direct measurements; in brackets: Standard Deviation

#### Supplementary Table 2

| STOCK/TRANSGENE | SOURCE | NUMBER |
|---|---|---|
| Drosophila, GMR-FLP (X) | Chang et al. (1995) | N/A |
| Drosophila, GMR-Gal4 (II) | Bloomington Drosophila Stock Center (BDSC) | 1104 |
| Drosophila, GMR-Gal4 (III) | BDSC | 29967 |
| Drosophila, FRT80B, tub-Gal80 | BDSC | 5191 |
| Drosophila, FRT82B, tub-Gal80 | BDSC | 5135 |
| Drosophila, FRT42D, GMR-Gal80 | This paper. GMR-Gal80: Gift from Thomas Clandinin. | N/A |
| Drosophila, FRT40A, tub-Gal80 | BDSC | 5192 |
| Drosophila, FRT2A, tub-Gal80 | BDSC | 5190 |
| Drosophila, FRT40A | BDSC | 8212 |
| Drosophila, FRT42D | BDSC | 1802 |
| Drosophila, FRT80B | BDSC | 8214 |
| Drosophila, FRT82B | BDSC | 5619 |
| Drosophila, FRT2A | BDSC | 1997 |
| Drosophila, FRT82B, *syd-1*$^{w46}$ | Holbrook et al. (2012) | N/A |
| Drosophila, FRT82B, *syd-1*$^{dRhoGAP}$ | This paper. | N/A |
| Drosophila, FRT40A, *liprin-α*$^{E}$ | Choe et al. (2006) | N/A |
| Drosophila, FRT40A, *lar*$^{2127}$ | Maurel-Zaffran et al. (2001) | N/A |
| Drosophila, FRT2A, *trio*$^{3}$ | Newsome et al. (2010) | 9130 |
| Drosophila, / UAS-Brp-RNAi$^{B3}$, UAS-Brp-RNAi$^{C8}$ | Wagh et al. (2006) | N/A |
| Drosophila, UAS-CD4-tdGFP (II) | BDSC | 35839 |
| Drosophila, UAS-CD4-tdGFP (III) | BDSC | 35836 |
| Drosophila, UAS-CD4-tdTomato (III) | BDSC | 35837 |
| Drosophila, UAS-BrpD3-GFP (III) | Schmid et al. (2008) | N/A |
| Drosophila, UAS-BrpD3-mKate2 (II and III) | This paper. | N/A |
| Drosophila, UAS-Liprinα-GFP | Fouquet et al. (2009) | N/A |
| Drosophila, UAS-GFP-Syd1 | Owald et al. (2010) | N/A |
| Drosophila, GMR-myr-tdTomato (II and III) | Gift from S.Lawrence Zipursky | N/A |
